# ROS induction as a strategy to target persister cancer cells with low metabolic activity in NRAS mutated melanoma

**DOI:** 10.1101/2022.10.19.512839

**Authors:** Ossia M. Eichhoff, Corinne I. Stoffel, Jan Käsler, Luzia Briker, Patrick Turko, Gergely Karsai, Nina Zila, Verena Paulitschke, Phil F. Cheng, Alexander Leitner, Andrea Bileck, Nicola Zamboni, Anja Irmisch, Zsolt Balazs, Aizhan Tastanova, Susana Pascoal, Pål Johansen, Rebekka Wegmann, Julien Mena, Alaa Othman, Vasanthi S. Viswanathan, Judith Wenzina, Andrea Aloia, Annalisa Saltari, Andreas Dzung, TuPro Consortium, Michael Krauthammer, Stuart L. Schreiber, Thorsten Hornemann, Martin Distel, Berend Snijder, Reinhard Dummer, Mitchell P. Levesque

## Abstract

Metabolic reprogramming is an emerging hallmark of resistance to cancer therapy but may generate vulnerabilities that can be targeted with small molecules. Multi-omics analysis revealed that NRAS-mutated melanoma cells with a mesenchymal transcriptional profile adopt a quiescent metabolic program to resist cellular stress response induced by MEK-inhibitor resistance. However, as a result of elevated baseline ROS levels, these cells become highly sensitive to ROS induction. *In vivo* xenograft experiments and single-cell RNA sequencing demonstrated that intra-tumor heterogeneity requires the combination of a ROS-inducer and a MEK-inhibitor to target both tumor growth and metastasis. By *ex vivo* pharmacoscopy of 62 human metastatic melanomas, we found that MEK-inhibitor resistant tumors significantly benefitted from the combination therapy.

Finally, we profiled 486 cancer cell lines and revealed that oxidative stress responses and translational suppression are biomarkers of ROS-inducer sensitivity, independent of cancer indication. These findings link transcriptional plasticity to a metabolic phenotype that can be inhibited by ROS-inducers in melanoma and other cancers.

**Statement of Significance:** Targeted-therapy resistance in cancer arises from genetic selection and both transcriptional and metabolic adaptation. We show that metabolic reprogramming sensitizes resistant cells to ROS-induction in combination with pathway inhibitors. Predictive biomarkers of metabolic sensitivity to ROS-inducing agents were identified in many cancer entities, highlighting the generalizability of this treatment approach.

**Graphical summary:** 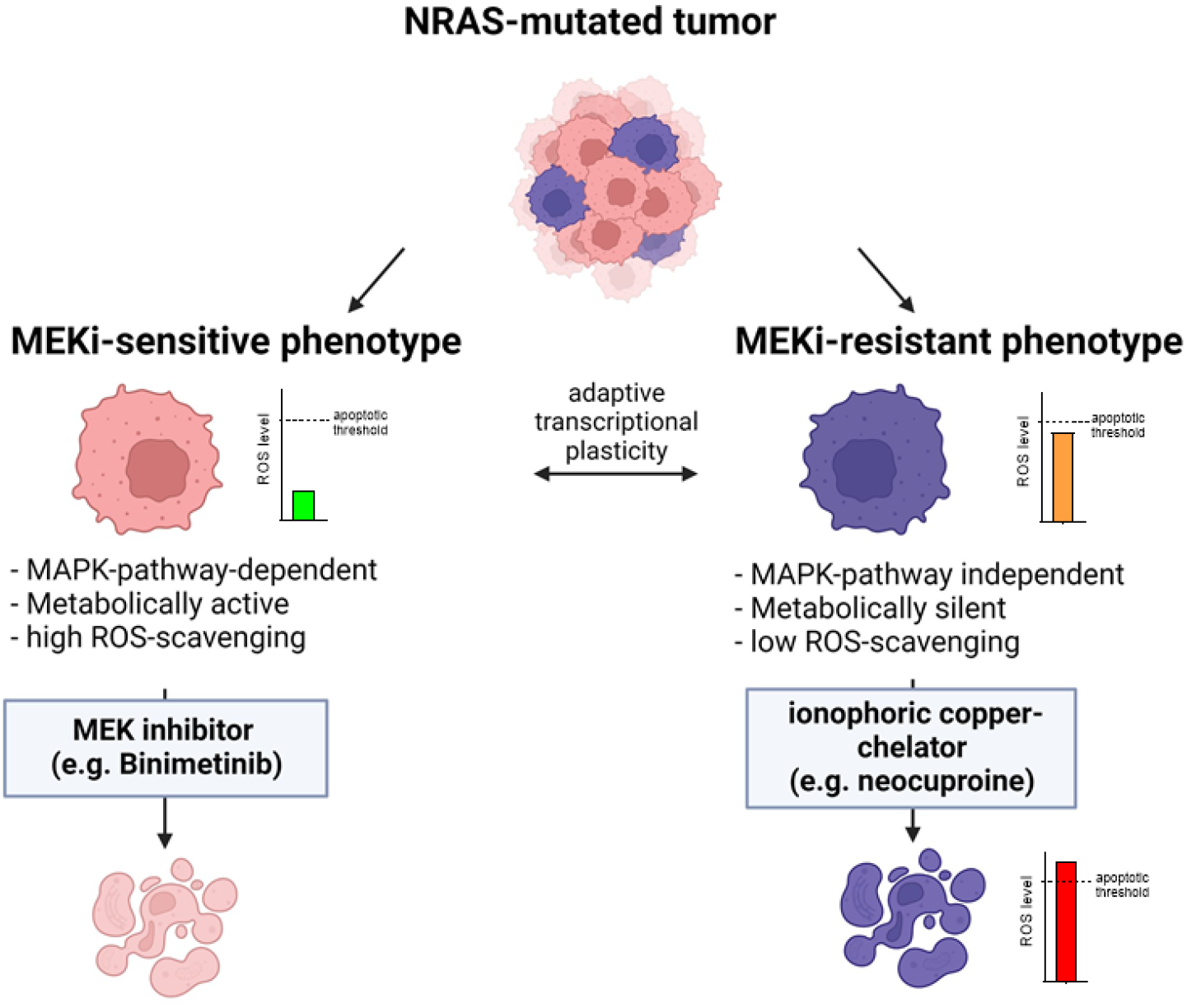

## Introduction

Activating mutations in the NRAS gene occur in approximately 20-30% of melanoma patients, resulting in hyperactivation of MAPK signaling and aggressive diseases [1, 2]. In recent years, targeting the MAPK pathway in BRAF-mutant melanoma has led to a dramatic improvement in the 5-year overall survival of BRAF-mutated patients [3]. However, intensive efforts to develop small molecules that directly bind to mutated NRAS proteins have failed [4]. Given the lack of targeted therapies for patients with NRAS-mutated melanoma, immunotherapy is the recommended first-line treatment [5]. MEK inhibition in NRAS-mutated melanoma can prolong progression-free survival and has been suggested as a treatment option after immunotherapy failure [6, 7].

Since targeting MAPK signaling in NRAS-mutated melanoma is only beneficial to a small subset of patients (response rate for binimetinib is 15-20%), it is unclear whether other vulnerabilities in these cells might be druggable, for which genome-wide screens have been conducted [8].

Cell metabolism is an important hallmark of cancer and has an impact on tumorigenesis and metastasis, leading to deregulation of the intracellular redox balance [9]. Consequently, alterations of intracellular ROS (reactive oxygen species) levels dictate cellular phenotypes and modulate oncogenic signaling. Elevated ROS levels have been demonstrated to promote carcinogenesis and cancer progression by amplifying genomic instability through DNA damage and directly stimulating cancer-promoting signaling pathways in tumor cells [10]. Oncogenic BRAF mutations and the resulting hyperactivation of MAPK signaling increase the glycolytic phenotype in melanoma, thereby delivering antioxidant defense mechanisms through the pentose phosphate pathway, which ensures cancer cell survival and proliferation by linking glycolysis to intracellular ROS control [11, 12]. In contrast to the Warburg effect, in which tumors produce energy (in the form of ATP (adenosine triphosphate)) through glycolysis, recent studies have shown that melanomas treated with MAPK pathway inhibitors have reduced glycolysis and elevated oxygen consumption levels, and therefore high oxidative phosphorylation rates (OXPHOS), resulting in elevated beta-oxidation and altered generation of superoxide anions [13–15]. A similar intensive analysis of the inhibition of the MAPK pathway and the resulting metabolic phenotype in NRAS-mutated melanoma is lacking. In non-cancer cells, ROS are produced at low concentrations and are therefore effectively scavenged by the potent cellular antioxidant system. Therefore, targeting ROS in aggressive cancer cells with elevated ROS levels and impaired ROS-scavenging systems is a promising treatment strategy [16].

In this study, we employed a high-throughput screening approach to target NRAS-mutated MEK inhibitor-resistant melanoma cells. This led to the discovery of neocuproine, which targets changes in the energy metabolism of NRAS-mutated melanoma cells by increasing the ROS levels. Because of the high sensitivity of NRAS-mutated, treatment-resistant, and mesenchymal melanoma cells to neocuproine, we were able to elucidate novel differences in the metabolism of MEK inhibitor-resistant melanoma cells and their connection to transcriptional cell states and phenotype plasticity. *Ex vivo* tumor analysis, xenograft responses, and broad cancer cell line panel screening point towards a general clinical potential of ROS inducers in the treatment of cancer as well as in combination with kinase inhibitors.

## Material & Methods

### Primary human melanoma cell lines and treatments

Patient-derived melanoma cell lines were provided by the Melanoma Biobank, University Hospital Zurich, which were established from surplus material from primary cutaneous and metastatic melanoma removed by surgery. Written informed consent was obtained from patients and the study was approved by the local IRB (EK647 and EK800; BASEC-Nr.2017-00494; BASEC-Nr.2014-0425). Melanoma cells were isolated from the tissue biopsies and grown as previously described [17]. Melanoma cell lines were maintained in RPMI 1640 (Gibco, Cat# R8758) supplemented with 5 mM glutamine (Gibco, Cat# 25030-081), 1 mM sodium pyruvate (Sigma-Aldrich, Cat# S8636), and 10% heat-inactivated fetal calf serum (FCS, Lonza, Switzerland) and cultured at 37°C and 5% CO2. Small-molecule kinase inhibitors: BRAFi (Selleckchem, Encorafenib, Cat#S7108); MEKi (Selleckchem, Binimetinib, Cat#S7007)) and all STO compounds were derived from the ActiTarg-K Kinase Modulators Library (TimTec, Delaware, USA). The other compounds used in this study were N-acetyl-L-cysteine (NAC, Sigma-Aldrich, Cat#A7250), dichloroacetate (DCA, Sigma-Aldrich, Cat#347795), and neocuproine (Sigma-Aldrich, Cat#N1501).

### Proliferation assay

Cell proliferation and IC50 values (half-inhibitory concentration) were determined using Resazurin reagent (Sigma-Aldrich, Cat#R7017). Briefly, melanoma cells were seeded in 96-well culture plates and allowed to adhere overnight at 37°C and 5% CO2. At different time points, the cells were treated with different concentrations of compounds for 72h. On the day of the assay, the cell culture medium was exchanged with 10% Resazurin solution (Resazurin stock solution:0.15 mg/ml in PBS (Gibco, Cat#10010-015)). Fluorescence was read using a plate reader (exc 535 nm, em 595 nm), and IC50 values were determined using GraphPad Prism software.

### High-throughput compound screening

Compound screening of 960 potential kinase inhibitors (ActiTarg-K Kinase Modulators Library, TimTec) was performed together with NEXUS Personalized Health Technologies (ETH Zurich, Switzerland). MEK inhibitor-sensitive and MEK inhibitor-resistant melanoma cell lines (three sensitive and three resistant) were seeded in 96-well plates and allowed to adhere overnight at 37°C and 5% CO2. On the day of the assay, 1 μM of each of the 960 compounds was added to each well of a 96-well plate and incubated for an additional 72 h. Cell proliferation was measured using a Resazurin solution, as described above. The fluorescence values were normalized to [0, 100] using positive and negative controls. Positive controls consisted of 10 μM Taxol (Paclitaxel, Selleckchem, Cat#S1150) or 10 μM ERK inhibitor (SCH772984, Selleckchem, Cat#S7101), which resulted in the lowest viability. Negative controls were treated with 0.1% DMSO. The normalized fluorescence was calculated as follows: (raw fluorescence – taxol/ERK) / (DMSO – ERK) × 100. Compounds that reduced the mean normalized viability to < 50% were targeted for further study.

### 3D-spheroid invasion assay

3D spheroids were obtained by growing melanoma cell lines under nonadhesive conditions. Briefly, 96-well plates were coated with 1.5% agar dissolved in RPMI medium and incubated under UV light in laminar flow for 30 min. 2^x^10^3^ cells/well were seeded on top of the agar and 3D sphere formation was observed 24-48 hours later under a light microscope. In experiments using line M160915, 3D spheroids were generated using ultra-low attachment surface spheroid microplates (Corning, Cat# 4520). 3D spheroids were collected carefully in 1.5 ml Eppendorf tubes and rested on ice. Per 3 ml of collagen-matrix 2.3 ml Collagen type I (from rat tails, Corning, Life Sciences, Cat# 354236) was mixed on ice with 0.3 ml DMEM base medium, 0.3 ml FCS, 25 μl glutamine, 60 μl sodium bicarbonate (7.5% solution, Sigma-Aldrich, Cat# S8761) and 30 μl antibiotic-antimycotic (Gibco, Cat# 15240-062). The access medium was carefully removed from the 3D spheres, and the spheres were mixed with 100 μl collagen-matrix and transferred to an agar-coated 96-well plate. 3D spheroids were incubated for 1 h at 37°C and 5% CO2 to polymerize the collagen matrix and overlaid with 100 μl of melanoma cell line medium. Once the first sign of cell invasion was visible under a light microscope (16-24 hours), the melanoma cell line medium was removed and replaced with compound-containing melanoma cell culture medium. Treatments were replenished once every 5 days, after which the cell culture medium was replaced with one containing 8 μM calcein-AM (Sigma-Aldrich, Cat# 17783) and ethidium homodimers (Sigma-Aldrich, Cat# 46043) and incubated for 1 h at 37°C and 5% CO_2_. The embedded 3D spheroids were imaged using a Leica DMi8 fluorescence microscope. Spheroid viability and invasion were analyzed using Adobe Photoshop (RRID:SCR_014199) and ImageJ (RRID:SCR_003070) Fsoftware, respectively. Viability was estimated by the ratio of the pixel intensity of calcein-AM staining and ethidium-homodimers. The area of invasion was calculated using the ImageJ software. All measurements were performed using at least three individual 3D spheroids.

### FACS analysis for the quantification of ROS and apoptosis

The amount of intracellular ROS was measured using 2’,7’-dichlorofluorescein diacetate (DCF-DA, Sigma-Aldrich, Cat# 35845). Briefly, cells were incubated for 30 min with fresh medium containing 1 μM DCF-DA (Sigma-Aldrich, CatNr. 287810) and analyzed using FACS. Mitochondrial ROS production (formation of superoxides) was detected using 5 μM MitoSox (Thermo Fischer, Cat# M36008) reagent according to the manufacturer’s protocol. To measure the activation of apoptosis, we used the Caspase-3/7 Detection Reagent (CellEvent™ Caspase-3/7 Green Detection Reagent, Thermo Fischer, Cat# C10423) at a concentration of 4 μM. Alternatively, the cells were stained with Annexin V/PI using the Apoptosis Detection Kit according to the manufacturer’s protocol (BioLegend, Cat# 640945). Cells were acquired on a BD LSR Fortessa, and data were analyzed using the FlowJo software (RRID:SCR_008520).

### Orthologic ex vivo slice cultures

Fresh tumor material was collected after surgery from consenting melanoma patients with confirmed NRAS mutations and kept in melanoma culture medium containing antibiotics (Gibco, Cat# 15140122) until the material was sectioned into 400 μm thick slices using a Leica microtome with a vibrating blade (VT 1200 S). Tumor slices were transfered onto 0.4 μm Millicell cell culture (Millipore, Cat# PICM03050) inserts and incubated with 1 ml of melanoma cell culture medium containing antibiotics for an additional 24 h when the medium was replaced with medium containing either vehicle, neocuproine (1 μM), MEK inhibitor binimetinib (0.5 μM), or the combination of neocuproine and binimetinib. The tumor slices were then placed on a nitrocellulose membrane to prevent folding, and the tissue was fixed in 4% buffered formalin. Slice cultures were subjected to immunohistochemistry, as described above (see section IHC).

### Immunohistochemistry

Fixed slice cultures were embedded in paraffin blocks, cut for immunohistochemical staining using the alkaline phosphatase-anti-alkaline phosphatase technique, and counterstained using hematoxylin (Leica, Bond Polymer Refine Red Detection Kit, Cat# DS9390; RRID:AB_2891238). The antibodies used were directed against S100 (Leica, Clone NCL-l-S100p, RRID:AB_442132, dilution 1:600), Ki67 (Dako, clone MIB-1, Cat# M7240, dilution 1:50), phospho-S6 Ribosomal protein (Ser240/244, D68F8, Cell Signaling #5364, RRID:AB_10694233, dilution 1:1000), phospho-p44/42 MAPK (Erk1/2) (Thr202/Tyr204, 20G11, Cell Signaling #4376, RRID:AB_331772, dilution 1:1000), Caveolin-1 (E249, Abcam ab32577, RRID:AB_725987), MelanA (Novus, NBP1-30151, RRID:AB_1987285, dilution 1:200) and .INHBA (Abcam, ab56057, RRID:AB_881196, dilution 1:100)

For further analysis, the slides were scanned using a slide scanner (Aperio) and ImageScope software (RRID:SCR_020993). Images were analyzed using QPath software (RRID:SCR_018257).

### Western Blotting

Cells were washed twice in ice-cold PBS, and proteins were solubilized in RIPA lysis buffer containing 20 mM Tris-HCl (pH 7.5), 1% Triton X-100, 150 mmol/L NaCl, 10% glycerol, and complete mini protease inhibitor (Roche Diagnostics Cat#11836170001). The protein concentration was measured using the Bio-Rad Dc Protein Assay (Bio-Rad, Cat#5000112) according to the manufacturer’s protocol. Proteins were separated on a NuPAGE 10% Bis-Tris gel (Invitrogen, Cat# NP0315) under denaturing and reducing conditions, followed by transfer onto a nitrocellulose membrane. Membranes were probed with anti-PARP (Cell Signaling Technology Cat# 9542, RRID:AB_2160739, dilution 1:1000), anti-pAMPK (Cell Signaling Technology Cat# 2535, RRID:AB_331250, dilution 1:1000), or anti-GAPDH (Cell Signaling Technology Cat# 2118, RRID:AB_561053, dilution 1:1000), followed by incubation with horseradish peroxidase-conjugated secondary antibodies (Cell Signaling Technology Cat# 7074, RRID:AB_2099233, dilution 1:1000). Bound antibodies were detected using chemiluminescence (ECL, GE Healthcare, Cat# GERPN2232) and imaged using the Fusion Fx Imaging System (Vilber, France).

### GSH assay

A glutathione assay bioassay kit (QuantiChrom, Cat# DIGT-250) was used to quantify reduced glutathione levels in melanoma cells, following the manufacturer’s instructions. 24 hours before performing the assay, the cells were provided with fresh medium. Briefly, 5 × 10^6^ cells were collected by centrifugation, washed in cold PBS, and lysed in 1 ml of cold lysis buffer containing PBS and 0.5% NP-40. All samples were collected in triplicate. The suspension was centrifuged, and the supernatant was mixed equally with Reagent A. The resulting turbidity was removed by centrifugation, and 100 μl of Reagent B was added to 200 μl of clear sample/Reagent A mixture. After 5 min, the optical density of the sample/reagent mixture was measured using a standard plate reader (Tecan Infinite 200 PRO). The GSH concentration of the samples was extrapolated from a GSH standard curve using GraphPad Prism software. All data represent the mean of at least three independent measurements.

### Zebrafish *in vivo* experiments

Zebrafish (Danio rerio) were maintained under standard conditions according to the guidelines of local authorities under licenses GZ208969/2015/18, GZ: 565304/2014/6, and GZ:534619/2014/4.

Tg(fli:GFP) embryos were raised at 28°C until 48 h post fertilization (hpf), dechorionated, anesthetized using 1x tricaine in E3 medium (0.16 g/l tricaine (Sigma-Aldrich Chemie GmbH, Germany), adjusted to pH 7 with 1M Tris pH 9.5, in E3), and placed in a bed of 2% agarose (Biozym LE Agarose, Vienna, Austria) in E3 medium on a petri dish lid for xenotransplantation. Xenotransplantation was performed using injection capillaries (glass capillaries GB100T-8P, without filament, Science Products GmbH, Germany) pulled with a needle puller (P-97, Sutter Instruments, Novato, USA) mounted on a micromanipulator (M3301R, World Precision Instruments Inc., Germany) and connected to a microinjector (Femto Jet 4i, Eppendorf, Germany).

Melanoma cells were maintained in culture medium containing 50 U/ml streptomycin/penicillin (Gibco, Thermo Fisher Scientific, Waltham, MA, USA) until they reached 90% confluency. DiI (D3911, Invitrogen, Thermo Fisher Scientific, Waltham, MA, USA) was diluted to 1:1000 in PBS and added to the cells on ice in the dark. After 30 min of incubation, the cells were detached and centrifuged for 5 min at 1200 rpm, followed by two washing steps with PBS without calcium and magnesium. Melanoma cells labelled with DiI were injected (appr. Two hundred cells/nl) into the perivitelline space (PVS) of zebrafish embryos at 48 hpf.

Immediately after injection, xenografted embryos were selected for successful transplantation of tumor cells and maintained at 35°C. Drug treatment with monotherapy MEKi binimetinib (500 nM), neocuproine (STO13881) (1 μM) and the combination binimetinib+neocuproine was applied from 24 hours post injection (hpi) till 96 hpi. DMSO was used as a control.

#### Imaging

At 96 hpi, xenografted zebrafish were anesthetized in 1x tricaine/E3, placed on the same side, and pictures were taken using an Axio Zoom.V16 fluorescence stereo zoom microscope with an Axiocam 503 color camera from Zeiss (Zeiss, Germany). Images were collected using Zeiss image software ZEN.

### Image-based ex vivo drug response testing on melanoma patient biopsies

#### Generation of drug assay plates

Between 1 and 100 nl of 5 mM drug stock (corresponding to final concentrations between 0.1 and 10 μM in the assay) and control (DMSO) are transferred to Corning 384-well, tissue-culture treated clear bottom plates by a Labcyte Echo liquid handler. Each drug plate contained multiple replicates for each drug and its concentration. Drugs are added either as single drugs or combinations of multiple drugs. Combinations were generated by transferring multiple compounds to the same well using an echo liquid handler. Once the plates were generated, they were stored at −20°C.

#### Sample processing

The single-cell suspension obtained from the TumorProfiler Central Lab was diluted in growth medium (RPMI1640 + GlutamaX (Gibco) + 10% FBS (Gibco) + 1% sodium pyruvate (Gibco) + 1% Anti-Anti (Gibco)). Around 2500-5000 cells per well were seeded into a 384-well plate containing a single drug or drug combination in each well. For each tested drug and drug combination, the plate contained at least three replicates at two different concentrations. After overnight incubation, cells were fixed with 10% formalin [Sigma]. Then, the cells were stained with DAPI [Sigma], a fluorescent dye used for nucleus detection, and a panel of melanoma-specific fluorescent antibodies, as described in detail in Table 1. Cells were imaged using an Opera Phenix automated microscope at 10x magnification, resulting in nine non-overlapping images per well.

**Table 1:** List of antibodies used to perform single-cell drug responses by pharmacoscopy

| Primary antibody | Short name | Note |
| --- | --- | --- |
| Rabbit monoclonal<br>[EPR14335] to SOX9<br>(Abcam Cat#<br>ab202517;Alexa Fluor<br>594) | Sox9 |  |
| *Rabbit monoclonal [EP1576Y] to S100 beta (Abcam Cat# ab196442, RRID:AB_2722596)* | *S100B* |  |
| *Mouse monoclonal to HMB45 (BioLegend Cat# 911506, RRID:AB_2750178)* | *HMB45* | *For samples M8BO8, MY5BB, M8WRI and MTPTA, the following combination of primary and secondary antibody were used instead of the primary conjugated:*<br>*HMB-45 Monoclonal (BioLegend Cat# 911501, RRID:AB_2565112)*<br>*Alexa Fluor® 647 anti-mouse IgG1, (BioLegend Cat# 406617, RRID:AB_2563476)* |
| *Alexa Fluor® 594 anti-MART-1 Antibody, (BioLegend Cat# 917906, RRID:AB_2750107)* | *MelanA* | *For samples M8BO8, MY5BB, M8WRI and MTPTA,, the following combination of primary and secondary antibody were used instead of the primary conjugated:*<br>*Purified anti-MART-1, Mouse IgG2b, (BioLegend Cat# 917901, RRID:AB_2565202)*<br>*Goat anti-Mouse IgG2b Cross-Adsorbed Secondary Antibody Alexa Fluor 594, Invitrogen A-21145* |

#### Data analysis

Raw images generated by Opera Phenix were first analyzed with CellProfiler (v.2.2.0) to detect the location of nuclei and extract fluorescence intensities for each single cell. After an initial quality filter that removes cell clumps and debris from the analysis, cell types were assigned by thresholding the fluorescence intensity. Melanoma cells are defined as those expressing any of the markers S100B, Sox9, MelanA, or HMB45; non-melanoma cells are thus negative for all markers. The melanoma-specific drug response score was calculated as follows:

For each well, the fraction of melanoma cells is calculated as

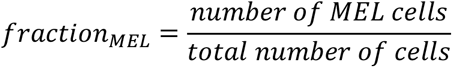

This value is then normalized to the fraction of melanoma cells in DMSO control wells, yielding a relative cancer cell fraction (RCF) per well:

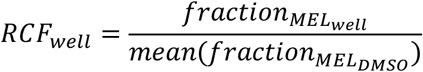

Finally, these values are subtracted from 1 and averaged across technical replicate wells and concentrations, yielding an area under the curve (AUC) for each drug:

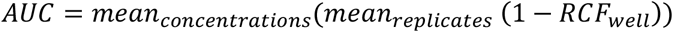

Treatments are ranked based on the AUC. An AUC > 0 indicates an on target effect, AUC = 0 means no effect or equal sensitivity of the melanoma and non-melanoma cells, and AUC < 0 corresponds to toxicity / higher sensitivity of the non-melanoma cells.

#### QC criteria

Each sample was assigned a QC score between 1 (very poor) and 5 (excellent). This score was estimated manually by considering cell viability, technical reproducibility, and the presence of autofluorescence and debris in the microscopy images. Samples with QC scores less than 2 (n=10) were excluded from the analysis. Furthermore, we excluded mucosal (n=5) and ocular (n=11) melanomas, leaving 92 samples analyzed in this study.

#### Integration of ex vivo response scores with CyTOF measurements

Only samples that were measured using both CyTOF and ex vivo drug response testing (n=72) were included in this analysis. Marker intensities measured by CyTOF were averaged across all tumor cells per sample. These average intensities were then compared between ex vivo responders (AUC > 0.02) and non-responders (AUC ≤ 0.02) using one t-test per marker, followed by multiple testing corrections across all tested markers using the Benjamini-Hochberg procedure.

### Melanoma Xenograft mouse model

All animal experiments were performed according to the protocols approved by the Swiss Federal Food Safety and Veterinary Office (license Nr. ZH/09517). We injected 500,000 cells of cell line M130227 or 10,000 cells of cell line M160915 together with Matrigel (Corning, Cat# CLS356234, dilution 1:1) into the flanks of BALB/c nude mice (Charles River, Germany). Treatment was started when tumors became palpable. Mice were then randomized into treatment groups and treated with either neocuproine (2 mg/kg in PBS) injected i.p. three times/week; binimetinib (MEK162, 30 mg/kg dissolved in 0.5% Tween-80 and 1% carboxymethylcellulose) was administered by oral gavage every working day (five times/week) or the combination of neocuproine and binimetinib. In the control group, mice were treated with a combination of vehicles alone. Xenograft experiments using a combination of binimetinib and elesclomol or binimetinib and disulfiram were carried out at EPO, Berlin, Germany. Here, 1 million cells (M130227) were injected together with Matrigel Corning, Cat# CLS356234, dilution 1:1) into the flanks of BALB/c nude mice. The mice were randomized into treatment groups when tumors were palpable and treated with binimetinib as described above. Elesclomol was injected three times per week (75 mg/kg in PBS). Tumor growth was monitored three times per week using calipers. The mice were euthanized when the tumor reached 1.5 cm a size. Draining lymph nodes were analyzed for micrometastasis using a TaqMan assay to distinguish between mouse and human DNA (for more details see also section in Material&Methods).

### Single-cell RNA Sequencing

Patient derived xenograft (PDX) tissue samples from mice were cryopreserved in 90% FBS (Biowest, Cat# S006420E01, batch no. S169419181H), and 10% DMSO (Cat. #102148154) in an Mr. Frosty cooling container (NALGENETM Cryo, Cat# 5100-0001). Before processing the samples for single-cell droplet generation, cryovials with slow frozen PDX tissue samples were quickly thawed in a water bath set to 37°C, re-suspended in 10 mL of ice cold RPMI 1640 with 1% BSA (Sigma-Aldrich, Cat# A7906,) and spun down at 300g for 5 min. Further, tissue samples were cut into small pieces and enzymatically digested for 30 min at 37 °C on a shaker with 5000U collagenase IV (Worthington, LS004188), 15 KU DNAse I (Sigma, Cat# D5025), 2 mL Accutase (Sigma, Cat # A6964) dissolved in RPMI with 2mM CaCl2 (Sigma-Aldrich, Cat# 746495,), and 0.5% BSA (Sigma-Aldrich, Cat# A7906,) 5 mL RPMI mixture. After incubation, the digested tissue was filtered through 100 μm nylon (Falcon, Cat# 352360) and then through 35-μm cell strainers (Falcon blue-capped FACS tubes, Cat# 352235). For samples with viability below 80%, apoptotic and dead cells were removed using immunomagnetic cell separation with the Annexin Dead Cell Removal Kit (StemCell Technologies, Cat# 17899) and EasySepTM Magnet (StemCell Technologies, Cat# 18000). If the cell pellet appeared red, red blood cell lysis was performed according to the manufacturer’s instructions (ROCHE, Cat# 11814389001). Cell number and viability were assessed on a Luna-FLTM Dual Fluorescence Cell counter (Logos Biosystems Inc.) using Photon slides (Ultra-low fluorescence counting slides, Logos Biosciences Inc., Cat# L12005) and acridine orange propidium iodide stain (Logos Biosciences Inc., AOPI, Cat# F23001), and optimal cell concentrations were set to 700-1100 cells/μL according to 10x Genomics protocols.

Single-cell droplets were generated using a 10x Genomics Chromium Single Cell Controller (10x Genomics, Cat# PN110211), Chromium Next GEM Single Cell 3 Kit v3.1 profiling kit (10x Genomics, Cat# PN-1000122), and Next Gem Chip G Single Cell Kit (10x Genomics, Cat# PN1000120). cDNA traces were amplified, and GEX libraries were constructed (10x Genomics Library construction kit, Cat# PN1000157; Single Index Kit T, Set A, Cat# PN1000213) according to the manufacturer’s instructions. A total of 6000 cells per sample were targeted, and the quality of cDNA traces and constructed gene expression libraries were evaluated on an Agilent 2100 Bioanalyzer (Agilent Technologies, system no. G1030AX) using a high-sensitivity DNA kit (Agilent Technologies, cat. no. 5067-4626). Sequencing strategy: Libraries were diluted to 10 nM and pooled at balanced ratios according to the target cell number. Paired-end sequencing (PE 28/8/0/91) was performed using an Illumina NovaSeq S2 flow cell. According to the 10x Genomics recommendation, 50,000 read pairs per cell were targeted for GEX coverage.

### TaqManPCR of lymph nodes for the examination of micrometastasis

After the termination of the *in vivo* experiments, tumor-draining mouse lymph nodes were collected, and DNA was extracted using the High Pure PCR template preparation kit (Roche, Cat# 11796828001) according to the manufacturer’s protocol. Total DNA was quantified for the amount of human and mouse DNA from the whole lymph node by using specific TaqMan probes, human or mouse, for the genomic region of PTGFR2, respectively [18]. The reaction was analyzed on a Via a7 qPCR machine (Applied Biosystems, USA).

### RNAseq

High-quality RNA was extracted using a QIAGEN RNeasy kit (Qiagen, Cat#74104). RNA capture was performed using TruSeq RNA Library Prep Kit v2 (Illumina, Cat#RS-122-2001; RS-122-2002). RNA sequencing was performed at a 150 bp single end on a HiSeq4000 at the Functional Genomic Center Zurich (FGZC). Gene-level counts were quantified from paired-end reads aligned to the GRCh38 genome using STAR (RRID:SCR_004463). A heat map was generated from ComplexHeatmap (RRID:SCR_017270) using R version 4.0.1 (http://www.R-project.org/). Phenotypes were assigned using signatures from Hoek [19].

### mRNA biomarker analysis (CCLE data)

The ranked list was made from the log odds of the CCLE analysis output. The MSigDB C2 collection was used for GSEA analysis [20, 21]. Network visualization of the pathways was performed with clusterProfiler (RRID:SCR_016884).

### Proteomics and GSEA

#### Preparation of cell pellets

Cells were washed twice and then harvested in ice-cold PBS (Gibco DPBS 1X, w/o calcium and magnesium). Samples were centrifuged at 4°C to generate cell pellets, supernatant was removed, and pellets were frozen immediately at −80°C for further analysis.

#### Tryptic digest of cell pellets

Cell pellets were thawed on ice and lysed using urea lysis buffer (8M urea lysis buffer, 5 mM EDTA disodium salt, 100 mM (NH4)HCO3, pH 8.0) with TCEP (10mM Tris(2-carboxyethyl)phosphine hydrochloride), protease (Roche cOmplete ULTRA Mini), and phosphatase (Roche PhosSTOP) inhibitors. After sonication and centrifugation, cell debris was discarded. The protein concentration was determined using a colorimetric assay (Pierce BCA Protein Assay Kit, Thermo Fisher Scientific) according to the manufacturer’s guidelines, with bovine serum albumin (BSA) as the standard. For proteomic analysis, we prepared in-solution digests using a variation of the FASP protocol [22], as previously described [23, 24]. Of each depleted serum and whole cell lysate 20 μg protein was concentrated onto a 10 kDa MWCO filter (molecular weight cut-off filter; Pall Nanosep Centrifugal Devices with Omega Membrane, #OD010). Concentrated proteins were reduced with 200 μl dithiothreitol (DTT) solution (5 mg/ml dissolved in 8 M guanidinium hydrochloride in 50 mM ammonium bicarbonate buffer, pH 8) and incubated at 56°C for 30 min. After centrifugation at 14,000 × g for 10 min, a washing step with 50 mM ammonium bicarbonate buffer was performed. For alkylation 200 μl iodoacetamide (IAA) solution (10 mg/ml in 8 M guanidinium hydrochloride in 50 mM ammonium bicarbonate buffer) was added and incubated at 30°C for 30 min in the dark. After centrifugation at 14,000 × g for 10 min, another washing step with 50 mM ammonium bicarbonate buffer was performed. Afterwards, filters were placed in a new Eppendorf tube, and 100 μl of 50 mM ammonium bicarbonate buffer as well as 10 μl of protease solution (Promega Trypsin/Lys-C Mix, Mass Spec Grade, #V5073, 0.1 μg/μl) were added, and incubated at 37°C for 18h. After digestion, peptide samples were eluted and collected peptides were acidified with 0.5% trifluoroacetic acid (TFA) to a final concentration of 1% TFA. A sample clean-up step with C-18 spin columns (Thermo Fisher Scientific Pierce C18 spin columns, #89870) was performed before peptide samples were finally dried at 40°C using a centrifugal vacuum concentrator (Labconco CentriVap) and stored at −20°C until MS analysis was performed.

#### LC-MS data acquisition

LC-MS/MS analysis was performed on an Easy nLC 1000 HPLC system coupled to an Orbitrap Elite mass spectrometer (Thermo Fisher Scientific), as previously described [24]. Peptides were separated on an Acclaim PepMap RSLC column (15 cm × 75 μm, Thermo Fisher Scientific) at a flow rate of 300 nl/min using a gradient from 5 to 30% mobile phase B for 180 min. The mobile phase compositions were A = water/acetonitrile/formic acid (98:2:0.15, v/v/v) and B = acetonitrile/water/formic acid (98:2:0.15, v/v/v). The Orbitrap Elite was operated in data-dependent acquisition mode with the detection of intact precursors in the orbitrap at a resolution of 120 000 and the detection of fragment ions in the linear ion trap at a normal scan speed. For each cycle, the 15 most abundant precursors with a charge state of +2 or higher were selected for fragmentation in the linear ion trap, with a normalized collision energy of 35%. Dynamic exclusion was performed for 30 s after 1 scan event.

#### LC-MS data analysis

Protein identification and label-free quantification (LFQ) were performed using the MaxQuant 1.5.2.8 software (RRID:SCR_014485), including the Andromeda search engine, and the Perseus statistical analysis package (RRID:SCR_015753) was used for subsequent statistical analysis [25]. Protein identification was achieved using the SwissProt database (version 102014 with 20 195 entries), a peptide tolerance of 25 ppm and a maximum of 2 missed cleavages. Furthermore, search criteria included a carbamidomethylation on cysteins as fixed modification and methionine oxidation as well as N-terminal protein acetylation as variable modifications and a minimum of two peptide identifications per protein, at least one of them unique. The match between runs option was used applying a 5 min match time window and a 15 min alignment time window. For both, peptides and proteins an FDR of less than 0.01 was applied. Before a two-sided t-test was performed, identified proteins were filtered for reversed sequences and common contaminants.

#### GSEA (gene set enrichment analysis)

Protein expression profiles were placed in a biological context using gene set enrichment analysis (GSEA). UniProt (UniProtKB, RRID:SCR_004426) accession numbers were mapped to gene symbols and GSEA was performed using the C2-Reactome Geneset collection (gsea-msigdb.org) with log2FC as a ranking metric, a minimum gene set size of 15, and a maximum gene set size of 500 [20]. An FDR of equal or below 0.25 was considered to be significant.

### Metabolic activity (Agilent Seahorse XF24 Flux analyzer)

To determine the extracellular acidification rate (ECAR) and oxygen consumption rate (OCR), melanoma cells were treated with DMSO or the indicated inhibitors for 48 h using an Agilent MitoStress Kit (Cat# 103015-100). Melanoma cells were seeded in quintuplicates on a Seahorse XF Microplate (Agilent, Cat# 100777-004) with melanoma culture medium supplemented with the corresponding drugs. Cells were incubated overnight in a humidified 37 °C incubator with 5% CO2. OCR and ECAR were measured using an XF24 extracellular flux analyzer (Seahorse Bioscience). Prior to performing the assay, the growth medium was exchanged with non-buffered XF Base DMEM medium (Agilent, Ca# 103334-100), supplemeted with 0.5 mM sodium pyruvate, 5.5 mM glucose (Sigma-Aldrich, Cat# G7021), 4 mM L-glutamine, and the appropriate compounds (binimetinib or neocuproine (STO13881)). plates were incubated for 1 h at 37 °C in a non-CO2 incubator. Three mitochondrial inhibitors (included in the MitoStress Kit), oligomycin (1 μM), FCCP (300 mM), and rotenone (1 μM), were sequentially injected after measurements according to the manufacturer’s standard protocol. OCR and ECAR were recorded, and values were normalized to the protein concentration in each well using a Bio-Rad Dc Protein Assay.

### Untargeted metabolomics

Melanoma cells (2 sensitive and 3 resistant cell lines) were seeded in 6-well plates at a density of 60-70% and allowed to adhere overnight. Subsequently, cells were treated with vehicle (0.1% DMSO) or neocuproine (STO13881) at increasing concentrations (0.5, 1, and 5 μM) in triplicate and incubated for an additional 24 h. Cells were rinsed with 75 mM ammonium carbonate (Cat Nr.379999, Sigma-Aldrich) at pH 7.4, and immediately frozen in liquid nitrogen. Metabolites were extracted by incubating cells with a 40:40:20 mixture of acetonitrile (Cat#271004, Sigma-Aldrich), methanol (Cat#106009, Millipore), and H20 for 10 min at −20°C. This procedure was repeated twice. Extracts were centrifuged at 4, °Cand the resulting supernatants were frozen at −80°C until analysis. Metabolite quantification was performed on an Agilent6550 QTOF instrument using flow injection analysis time-of-flight mass spectrometry. Detectable ions were putatively annotated by matching measured mass-to-charge ratios with theoretical masses of compounds listed in the human metabolome database v3.0 (http://www.hmdb.ca/ (HMDB, RRID:SCR_007712)) using a tolerance of 0.001 amu. Differentially abundant metabolites were categorized using the small-molecule pathway database (http://smpdb.ca/). To analyze the activation of the pentose phosphate pathway, melanoma cells were incubated in melanoma cell culture medium supplemented with 13C-labled glucose (200 mM; Sigma-Aldrich, CatNr. 389374).

### Lipidomics

Melanoma cells were grown to sub-confluence and fed fresh culture medium 24 h prior to harvesting. Three biological replicates of five Miollion cells were collected per cell line, and cell pellets were frozen at −80°C until analysis. Lipid extraction was performed as previously described [26] with some modifications. To 20 μL of sample, 1 ml of a mixture of methanol: MTBE: chloroform (MMC) 1.33:1:1 (v/v/v) was added. The MMC was fortified with the SPLASH mix of internal standards (Avanti Lipids) and 100 pmoles/ml of the following internal standards: d7-sphinganine (d18:0), d7-sphingosine (d18:1), dihydroceramide (d18:0:12:0), ceramide (d18:1/12:0), glucosylceramide (d18:1/8:0), sphingomyelin (18:1/12:0), and 50 pmoles/ml d7-sphingosine-1-phosphate. After brief vortexing, the samples were continuously mixed using a thermomixer (Eppendorf) at 25 °C (950 rpm, 30 min). Protein precipitate was obtained after centrifugation for 10 min at 16000 g at 25 °C. The single-phase supernatant was collected, dried under N2, and stored at −20 °C until analysis. Before analysis, the dried lipids were redissolved in 100μL of MeOH: isopropanol (1:1).

Liquid chromatography was performed as previously described [27] with some modifications. Lipids were separated by C30 reverse-phase chromatography. A Transcend TLX eluting pump (Thermo Scientific) was used with the following mobile phases: A) acetonitrile: water (6:4) with10mM ammonium acetate and 0.1% formic acid, and B) isopropanol: acetonitrile (9:1) with 10mM ammonium acetate and 0.1% formic acid. The C30 Accucore LC column (Thermo Scientific) with the dimensions 150 mm * 2.1 mm * 2.6μm (length*internal diameter*particle diameter) was used. The following gradient was used with a flow rate of 0.26 ml/minutes; 0.0-0.5 minutes (isocratic 30%B), 0.5-2 minutes (ramp 30-43% B), 2.10-12.0 minutes (ramp 43-55%B),12.0-18.0 minutes (ramp 65-85%),18.0-20.0 minutes (ramp 85%-100%B), 20-35 minutes (isocratic 100%B), 35-35.5 minutes (ramp 100-30% B) and 35.5-40 minutes (isocratic 30%B).

The liquid chromatography was coupled to a hybrid quadrupole-orbitrap mass spectrometer (Q-Exactive, Thermo Scientific). Data-dependent acquisition with positive and negative polarity switching is used. A full scan was used scanning from 220-3000m/z at a resolution of 70000 and AGC Target 3e6, while data-dependent scans (top10) were acquired using normalized collision energies (NCE) of 25, 30 and a resolution of 17,500 and AGC target of 1e5. Lipid identification was achieved using four criteria: 1) high accuracy and resolution with an accuracy within m/z within a 5 ppm shift from the predicted mass and a resolving power of 70000 at 200 m/z. 2) Isotopic pattern fitting to expected isotopic distribution. 3) Comparing the expected retention time to an in-house database, and 4) matching the fragmentation pattern to an in-house experimentally validated lipid fragmentation database. Quantification was done using single point calibration by comparing the area under the peak of each ceramide species to the area under the peak of the internal standard. Quality controls using a mixture of all samples were used at four concentration (1x,0.5x,0.25xand 0.125 ×). Triplicates of the QCs were measured, and the CV% for each of the lipids reported was below 20%. The following classes were identified in the current study: aclycarnitites, phospholipids (phosphatidylcholines, phosphatidylethanolamines, phosphatidylinositols,phosphatidylglycerols, and phosphatidyserines), sphingolipids (sphingoid base phosphates, ceramides, deoxyceramides, monohexosylceramides, and sphinogmeylins), and glycerolipids(diacylglycerol, triacylglycerol). Mass spectrometric data analysis was performed using Treacefinder software 4.1 (ThermoScientific) for peak picking, annotation, and matching to the in-house fragmentation database.

### Statistical analysis

All experiments in this study were performed with at least three replicates. Data analysis, unless described differently, was performed using Student’s t-test or 2-way ANOVA depending on the data format, using the GraphPad Prism software (RRID:SCR_002798). Statistical analysis of in vivo mouse survival data was performed using SPSS software (RRID:SCR_002865).

## Results

### High-throughput compound screen identified STO13881 (Neocuproine) as an inhibitor of NRAS-mutated and MEK-inhibitor resistant melanomas

Since treatment options are limited for NRAS-mutated melanoma patients, we performed high-throughput small-molecule compound screening to identify molecules that target NRAS-mutated and MEK inhibitor-resistant melanoma cells. We used patient-derived melanoma cell lines with confirmed NRAS mutations, including those which showed resistance to the MEK inhibitor binimetinib ((n=6; sensitive (M130515, M130429, M130427) resistant (M131205, M130219, M130227), suppl. Table ST1+2). Only four of the 960 compounds significantly inhibited the growth of melanoma cells by more than 50% on average across our cell panel at a concentration of 1 μM (Figure 1a, Suppl. Figure S1a). Interestingly, two compounds (STO13881 and STO12435) reduced the growth of NRAS-mutated MEK inhibitor (binimetinib)-resistant cells more compared to sensitive once (Suppl. Figure S1a). All four compounds were manually validated on the same NRAS-mutated melanoma cell panel (n=6) with dose-escalating concentrations to determine the precise IC50 values of each compound for the growth rate of the individual cell lines (Suppl. Figure S1 c-d). Only compound STO13881 significantly impaired cell growth more in MEK inhibitor-resistant cell lines (Figure 1 b-c). We confirmed STO13881 and neocuproine to have overlapping HPLC profiles (Suppl. Figure S2). We used patient-derived primary cell cultures of fibroblasts, melanocytes, and keratinocytes to show that heathy cells were not affected by neocuproine treatment at similar concentrations (Suppl. Figure S1 b). We confirmed that neocuproine targets resistant cell lines in an additional ten patient-derived melanoma lines with confirmed NRAS mutations (Suppl. Table ST1, Suppl. Table ST2). We observed significant differences in the sensitivity to neocuproine between MEKi-sensitive and resistant melanoma lines (hereafter referred to as sensitive and resistant, respectively; (Suppl. Figure S3 a-b)). To assess the additional benefit of neocuproine treatment and account for differences in sensitivity to MEKi, we calculated the IC50 ratio of neocuproine and MEKi (binimetinib) (Suppl. Table ST1). We plotted the neocuproine/MEKi ratios and observed significant separation between the resistant and sensitive cell lines (Figure 1 d). Next, we wanted to confirm our findings on resistant melanomas with NRAS mutations in a physiological cell culture model reflecting key features of solid tumors like cell-cell contacts as well as a microenvironment. We performed 3D spheroid invasion assays in a collagen type I matrix and examined the viability and area of invasion of 3D spheroids derived from the resistant and NRAS mutated melanoma cell line M160915 (Figures 1 e, S3 c-d). We found that neocuproine and its combination with MEKi had a significant effect on cell viability and invasion area. We recently reported that BRAF-mutated melanomas can acquire MAPK pathway resistance through additional NRAS mutations in the same cells [28]. We also investigated whether these cells were sensitive to neocuproine. 3D spheroids derived from M121224 (NRAS/BRAF-mutated) cells were embedded in a collagen I matrix and treated with BRAFi (i.e., encorafenib), MEKi, neocuproine, or their combinations. This patient progressed under encorafenib treatment, and we found that the co-occurrence of an additional NRAS mutation during treatment was associated with resistance to BRAFi. Consistent with this observation, spheroids thrived in the presence of BRAFi (Suppl. Figure S3 e-f). Interestingly, although the MEKi can target these cells in vitro, we found that 3D spheroids were still viable and invaded the collagen matrix under MEKi treatment. However, the viability and invasiveness of 3D spheroids was abrogated by neocuproine and the addition of MAPKi significantly enhanced this effect in a concentration-dependent manner.

**Figure 1:**
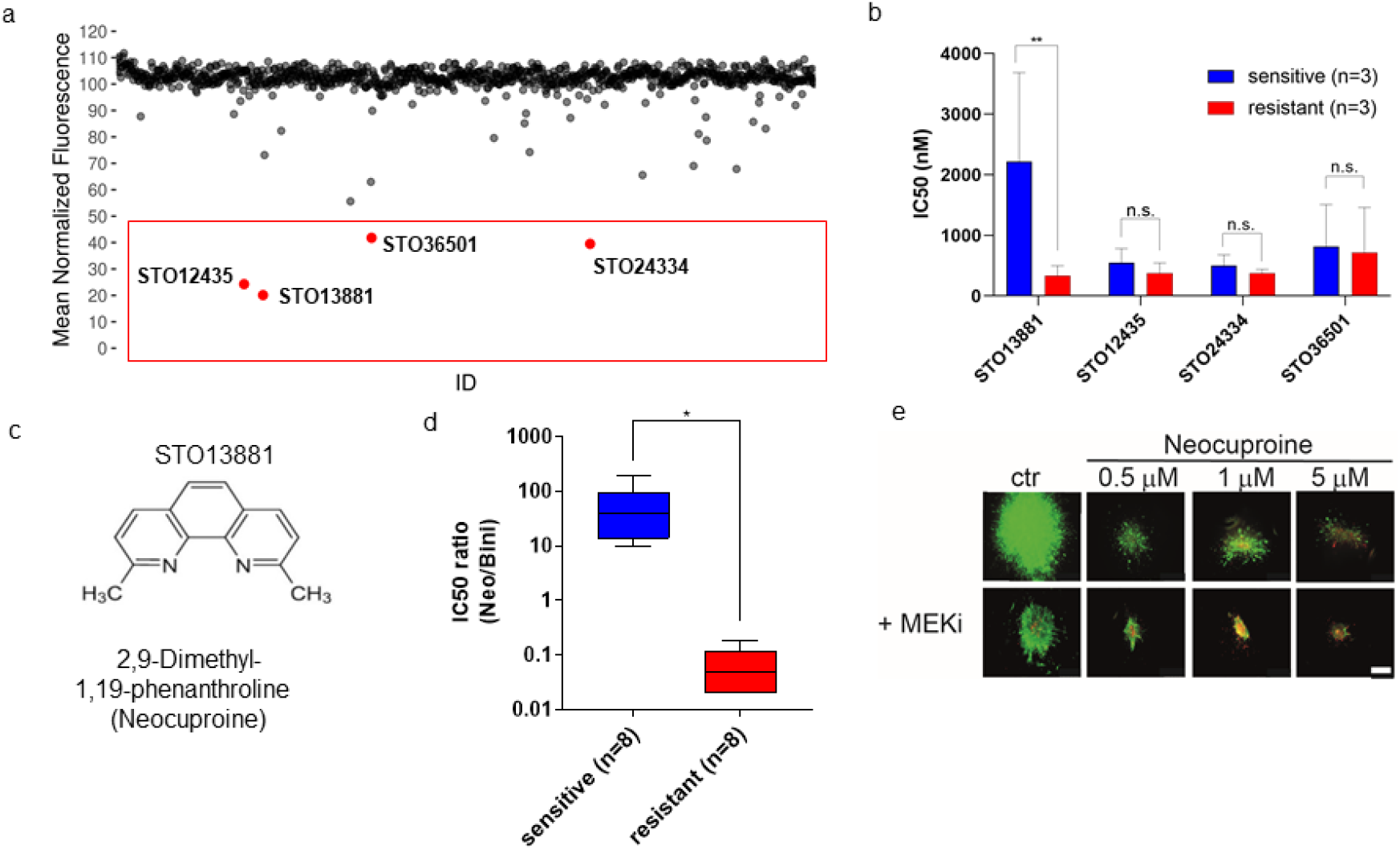
High-throughput compound screen revealed Neocuproine as a specific inhibitor for NRAS-mutated and resistant melanomas. (a) Results of the high-throughput screening (HTS) on a panel of 3 MEKi-sensitive (=sensitive) and 3 MEKi-resistant (=resistant) NRAS-mutated primary human-derived melanoma cell lines. Every dot represents the average and normalized growth (% to control) across cell lines per compound. (b) The bar graphs summarizes the IC50 values (in nM) of growth inhibition for sensitive versus resistant melanoma cell lines (3 sensitive vs 3 resistant) of all four compound hits (STO13881 p=0.01; two-way ANOVA). (c) Structural formula and the chemical name of compound STO13881 (=Neocuproine). (d) Neocuproine’s cytotoxicity was validated on a panel of 16 NRAS mutated cell li (8 sensitive vs 8 resistant). The box plots represent the summary of the IC50 ratios between Neocuproine (=Neo) and MEKi (Binimetinib) (p=0.01, Student’s t-test). (e) Fluorescent imaging of 3D melanoma spheroids embedded into a Collagen I matrix (scale bar=500 μM).

### Neocuproine impairs cell growth through ROS-induction

Neocuproine is known as a copper-chelating agent, which is rapidly reduced to neocuproine-Cu (I) complexes upon binding of Cu (II) [29]. In contrast to copper-chelating agents neocuproine is a “copper ionophore” which transports Cu from the extracellular space through cell membranes. The activity of ionophores is strictly Cu-dependent, and Cu ionophores are inactive under Cu-depleted conditions. Therefore, we used the copper-chelator trientine to deplete copper from the cell culture medium and reversed the effects of neocuproine on melanoma cells (Suppl. Figure S4 a). Although neocuproine-Cu complexes are stable, it has been observed that neocuproine-Cu (I) in combination with glutathione (GSH) can induce DNA breaks through oxidative mechanisms. Antioxidants, such as N-acetyl-L-cysteine (NAC), are ROS scavengers that reverse their cellular effects [30]. We found that the growth-inhibitory effects of neocuproine could be rescued by the antioxidant NAC, suggesting that this effect is caused by oxidative stress (Figure 2 a). Consistently, in 3D spheroid assays, collagen matrix invasion was significantly rescued by NAC co-treatment (Figure 2 b, Suppl. Figure S4 b-d). An early cellular response to DNA damage involves accumulation of γH2AX at DNA strand breaks induced by UV exposure, metabolic stress, and ROS [31]. Consistent with the observed upregulation of oxidative stress, we found a significant accumulation of γH2AX only in resistant melanoma cells treated with escalating doses of neocuproine (Figure 2 c). The observed DNA damage was also accompanied by an increase in resistant cells in the G2-phase in resistant cells (Suppl. Figure S4 e). It has been demonstrated that neocuproine-Cu complexes decrease mitochondrial membrane potential, which results in increased ROS production in astrocytes [32]. We found that, while cytoplasmic ROS were significantly upregulated in resistant melanoma, differences in superoxide production at the mitochondrial cell membranes were not detected (Suppl. FigureS5a). We observed a significant upregulation of cytoplasmic ROS upon neocuproine treatment only in resistant cells, which were sensitive to neocuproine (Figure 2 d). This effect was rescued by NAC, confirming the induction of ROS by neocuproine. Increasing evidence points towards the carcinogenic role of intracellular redox imbalance and aberrant ROS levels, which is consistent with the observed higher baseline ROS levels in our resistant melanoma lines [33]. In contrast, high levels of ROS are thought to induce cell death by inducing apoptosis [34]. Therefore, we investigated the induction of apoptosis by neocuproine. We observed increased caspase 3/7 activity, as well as an increased fraction of apoptotic cells in resistant melanoma cultures (Figures 2 e, Suppl. Figure S5 b-c). Consistent with these observations, we detected increased cleavage of PARP when resistant cells were treated with neocuproine (Figure S5 d). However, in the NRAS/BRAF double-mutated melanoma cell lines (i.e., M12224), we detected decreased levels of apoptosis as measured by caspase 3/7 activation compared to the NRAS-mutated and resistant cultures, suggesting that there are other mechanisms involved in cell death compared to cells that only harbor NRAS mutations. Thus, resistant cells experienced high basal ROS levels, which were further elevated by neocuproine to induce cell death via apoptosis. We then tested the hypothesis that cell lines resistant to MEKi are associated with high basal ROS levels, resulting in a resistant cell phenotype that corresponds to neocuproine sensitivity. Therefore, we analyzed our melanoma cell lines for basal ROS levels and responsiveness to neocuproine and MEKi. Basal ROS levels significantly correlated with MEKi inhibitor sensitivity (p<0.001, R-square 0.8) (Suppl. Figure S6 a). Applying MEKi-sensitivity as a categorical variable according to IC50 (i.e., whether a cell line was considered sensitive or resistant based on an IC50 threshold), we found a significant correlation with basal ROS levels (p<0.001, Suppl. Table ST1, Suppl. Figure S6 b). Finally, we tested for a correlation between the MEKi phenotype and neocuproine responsiveness (Figure 2 f) and found a strong relationship between basal ROS levels, inhibitor sensitivity, and responsiveness to neocuproine in NRAS-mutated melanoma (p<0.001).

**Figure 2:**
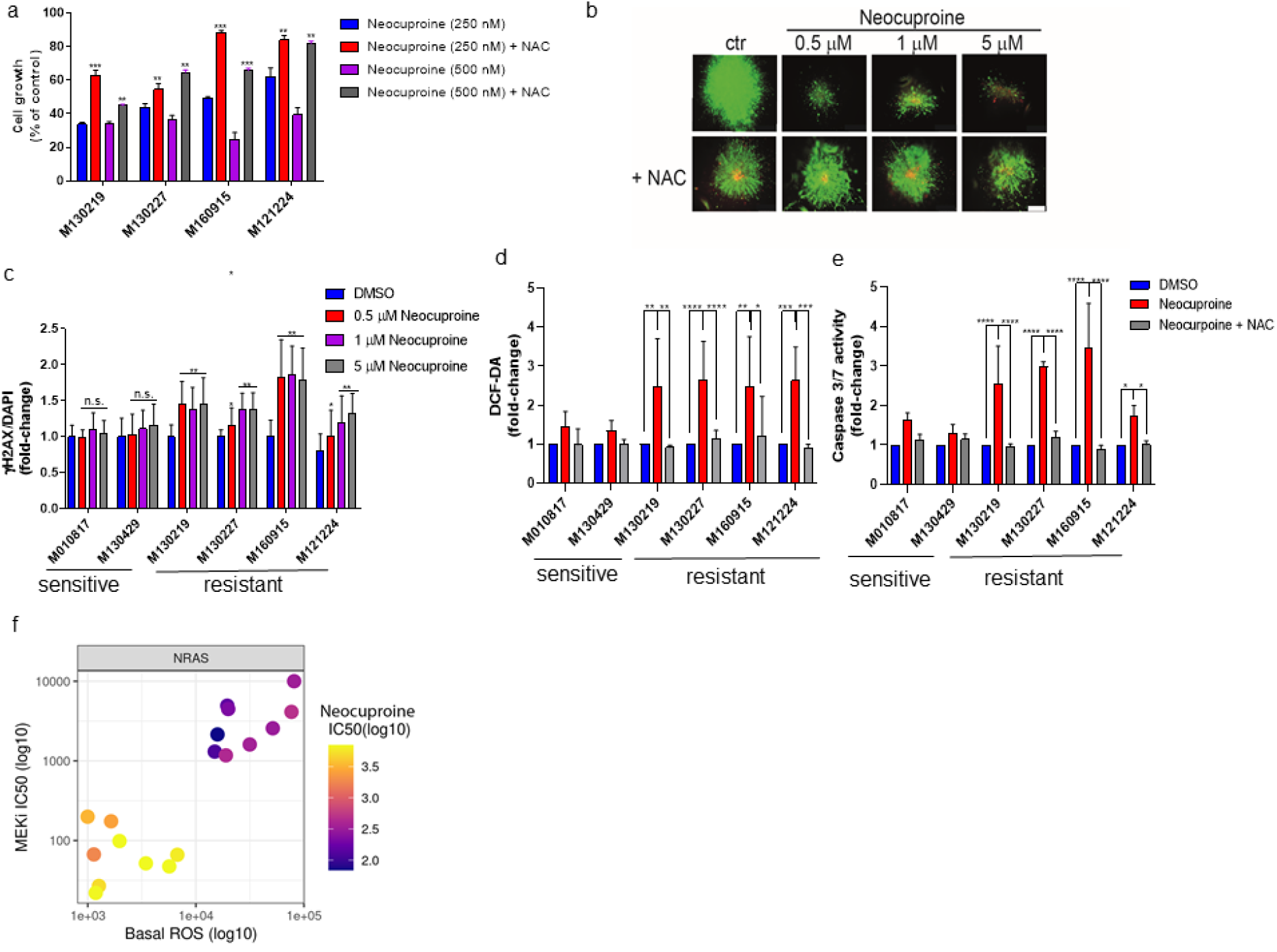
Neocuproine impairs cell growth through ROS-induction. (a) Growth inhibitory effects of the combination of Neocuproine and NAC on resistant melanomas. Cell lines were exposed to drugs for 72h and values are averages of replicates expressed relative to cell growth values in vehicle treated cells normalized to 100% (***p<0.001; **p<0.01, two-way ANOVA). (b) Fluorescent imaging of 3D spheroid invasion into a collagen type I matrix after the treatment with Neocuproine or in combination with NAC (scale bar=500 μM). (c) DNA double strand breaks were evaluated using γH2AX fluorescent staining in the presence of raising concentrations of Neocuproine (**p<0.01, ***p<0.001, two-way ANOVA). (d) Melanoma cells were treated with 1 μM of Neocuproine +/- NAC and analyzed by FACS with fluorescent probes DCF-DA. (**p<0.01, ***p<0.001, ****p<0.0001, two-way ANOVA) (e) Induction of apoptosis after the treatment with 1 μM Neocuproine +/- NAC was detected by FACS analysis using Caspase 3/7 activation assay (*p<0.05; ***p<0.001; ****p<0.0001; two-way ANOVA). (f) Correlation blot between MEKi, Neocuproine and basal ROS

### Sensitivity to neocuproine is associated with a mesenchymal cell phenotype and low metabolic activity

The “phenotype switching” model proposes that melanoma cells can toggle their transcriptional profiles in a dynamic process with similarities to the model of epithelial-mesenchymal transition (EMT) described for epithelial cancers [35, 36]. Genetic alterations are thought to modulate the stability and strength of the cellular phenotype and the threshold level of phenotype switching but may not drive transcriptional intra-tumoral phenotype switching [37, 38]. Moreover, melanoma phenotypes influence sensitivity to MAPK inhibitors [39]. We performed unsupervised hierarchical clustering on bulk RNAseq data from our NRAS mutated melanoma cohort and observed a splitting into the original melanoma cell phenotypes (“proliferative,” “intermediate” and “invasive”) described by Hoek et al. (Figure 3 a, Suppl Table ST3) [40, 41]. Cell lines with a mostly “proliferative” phenotype are sensitive to MEKi, but resistant to neocuproine. In contrast, we could associate most MEKi resistant cell lines with the “invasive or intermediate phenotype” described by Hoek et al, which we now call the “mesenchymal” state [42]. In recent years, considerable evidence has demonstrated that melanoma cells in this mesenchymal state have lost melanocytic differentiation markers and adapted EMT-like signatures attributed to drug tolerance [42–44]. Proteomics-based technologies have been recognized as powerful tools for understanding cell heterogeneity and resistance mechanisms in melanomas [24]. To understand the driving forces of our two cell phenotypes (sensitive vs. resistant to MEKi), we applied proteomics followed by gene set enrichment analysis (GSEA) to compare the pathway activity of sensitive and resistant melanoma (Figure 3 b, Suppl. Table S4 and S5). We performed differential protein expression analysis and hierarchical clustering of the proteomic data and ranked the 15 most informative proteins for sensitive and resistant melanoma cell lines. Consistent with our previously published transcriptional signatures of proliferative and invasive melanoma phenotypes by Hoek et al., we found that sensitive melanoma cells upregulated Mitf-driven and melanocyte-specific factors that are also involved in pigmentation (PMEL and MART1) or factors involved in the trafficking of melanocytic-associated factors (RAB38 and RAB32). Additionally, we identified several upregulated proteins that play roles in metabolism and energy production (i.e., UPP1, ABCD1, GALE, SLC7A5, and IDH1). Consequently, we observed downregulation of Mitf, Mitf-targets, and melanocytic protein expression in resistant melanomas, but upregulation of EMT-related proteins (e.g., TGM2, NT5E, SPARC, PLAT, and PRNP), proteins involved in drug resistance (IGFBP1 and PTRF), or those antagonizing glycolysis like PRKCDBP (Cav3), which is associated with lipid rafts, and loss of this factor induces Warburg metabolism and lactate production [45, 46]. GSEA revealed that melanoma cells sensitive to the melanocytic phenotype exhibited upregulated pathways involved in RNA transcription, MAPK pathway activation, and glucose metabolism. Resistant and mesenchymal melanoma cells down-regulate the MAPK pathway and glycolytic metabolism while upregulating two distinct pathways known to drive metastatic cancer cell phenotype and impact on EMT: “extracellular matrix organization” and “cell junction organization” [47, 48]. After treatment with neocuproine, we found only one pathway enriched in the sensitive (and neocuproine-resistant) cell cohort: “packaging of telomere ends” (Figure 3 b). “Packaging of telomere ends” includes mostly histone cluster 1 H4 upregulation, which might protect the DNA from oxidative stress. We did not identify any factors that were significantly associated with neocuproine responsiveness in the resistant cell lines.

**Figure 3:**
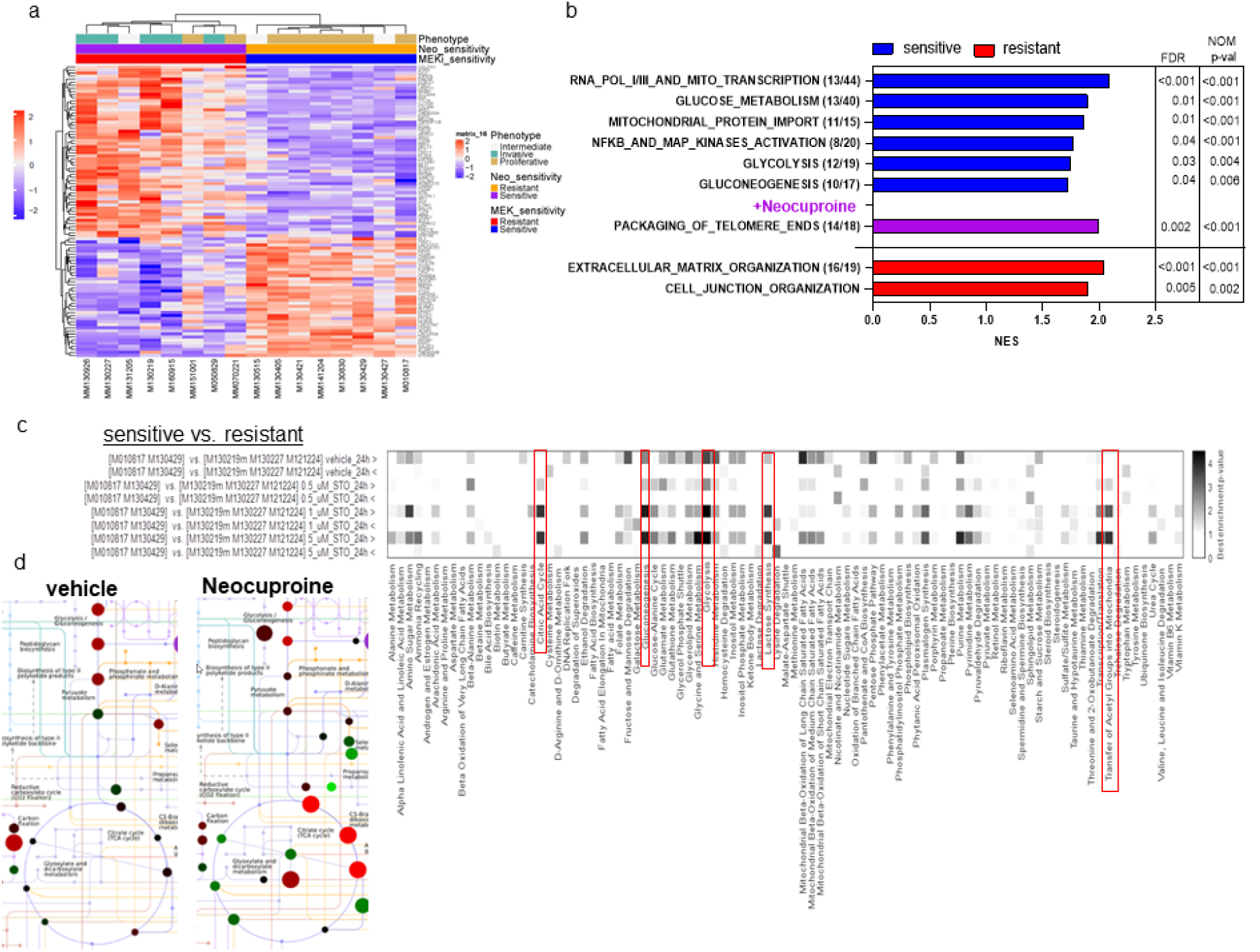
Sensitivity to Neocuproine is associated with a mesenchymal cell phenotype and low metabolic activity. (a) RNAseq data were subjected to unsupervised hierarchical clustering and transcripts up- or downregulated genes were plotted on a heatmap. Drug-sensitvity and melanoma cell phenotype were mapped to the cell lines (b) Proteomics data of sensitive and resistant melanomas also treated with 1 μM Neocuproine were analyzed with Reactome algorithm and only FDR significant pathways (cut-off: p<0.05) are shown as bar graphs (NES=normalized enrichment score (log2)). (c) Analysis of cell metabolites of sensitive and resistant melanomas including treatment with raising concentrations of Neocuproine (=STO) (0.5; 1; 5 μM for 24 h (hours)) were analyzed and pathway enrichment was plotted on a heatmap (see also Suppl Figure S6). > and < are indicating the direction of enrichments (d) Metabolite accumulation in the glycolytic and TCA pathway was visualized on a traffic map.

The enrichment of pathways involved in glycolytic processes in sensitive cell phenotypes prompted us to further investigate metabolic differences between sensitive and resistant melanomas. It is well known that metabolic pathways like glycolysis drive cellular detoxification by recycling of oxidized glutathione through NADPH [49]. Therefore, we performed mass spectrometry-based metabolomics on cell lysates from our cohort of NRAS-mutated melanomas. We confirmed enrichment of metabolites belonging to “Glycolysis,” “Gluconeogenesis,” “Phosphate pathway (PPP)” as well as “Mitochondrial beta-oxidation of long chain saturated fatty acids” (Figure 3 c, Figure S6 c). We did not find any metabolic pathways to be significantly enriched in resistant cell lines compared to sensitive ones, suggesting that resistance to MEKi is associated with a “silent” metabolic phenotype. Mass spectrometry-based lipidomic analysis of our cell cohorts revealed the separation of sensitive versus resistant cells based on their lipid profile (Suppl. Figure S7 a). Consistent with the metabolomic analysis, most of the significantly altered lipids were downregulated (75/89) in resistant cells (Suppl. Figure S7 b, Suppl. Table ST6). Most of these lipids are involved in energy metabolism (in red: Acylcarnitines, FAs, DAGs, and TAGs) and others essential for keratinocyte differentiation and phenotype switching (in green: Ceramides, Sphingomyelins, and Hexosylceramides) [50]. These data further support the “silent” metabolism and undifferentiated state of resistant melanoma cells already observed via metabolomics.

### Low Glycolysis and PPP activity sensitizes resistant melanoma cells to ROS

Metabolomics revealed that treatment of sensitive cell lines with 1 μM neocuproine for 24 hours further increased the metabolites related to glycolysis, glycolysis-related metabolic pathways (e.g., “Gluconeogenesis,” “Lactate synthesis,” “Pentose phosphate pathway”), and “Transfer of Acetyl Groups into Mitochondria,” Citric acid cycle” and “Plasminogen Synthesis” (Figure 3 c, Suppl. Figure S6 b). Therefore, we hypothesized that the defense against ROS induction generated by neocuproine is achieved through antioxidant scavenging by metabolic pathways, some of which are known to recycle GSH through NADPH (e.g. PPP) (Figure 3 d). To determine glycolytic cell capacity and the level of oxygen consumption, we employed Agilent Seahorse technology. We found that at baseline, sensitive melanoma cells (M130429 and M010817) had a significantly higher metabolic turnover rate determined by the extracellular acidification rate (ECAR, p<0.001), as well as a significantly higher oxygen consumption rate (OCR, p<0.001), in contrast to resistant melanoma cells (n=4; M130219, M130227, M160915, M121224) (Figure 4 a-b). To confirm the metabolomics data, which revealed a strong upregulation of glycolysis after the addition of neocuproine to sensitive melanoma cells, we measured the induction of ECAR during treatment. In response to neocuproine, sensitive melanoma cells significantly upregulated their glycolytic rate (measured as glycolytic reserve), whereas resistant cells did not, but in contrast, significantly decreased their glycolytic rate. Treatment of resistance cells with MEKi, significantly downregulated their glycolytic rate, suggesting strong regulation of glycolytic pathways through MAPK signaling in NRAS-mutated melanomas with melanocytic phenotypes (Figure 4 c-d). Upregulation of glycolytic pathways (e.g PPP) through intact MAPK signaling can supply sensitive melanoma cells with antioxidant defense metabolites (GSH and NADPH) through which they are protected from neocuproine-induced ROS. As a proof of concept, we treated sensitive melanomas with a combination of neocuproine and dichloroacetate (DCA), known to inhibit pyruvate dehydrogenase kinase and PPP [51]. We observed a significant decrease of neocurpoine’s IC50 in the presence of DCA (Figure 4 e). We also measured cytoplasmic ROS levels with DCF-DA and found a significant induction of ROS in sensitive cells treated with the combination of neocuproine and DCA, suggesting that inhibition of glycolysis/PPP is sufficient to sensitize cells to elevated ROS levels (Figure 4 f). DCA was used in a concentration of 10 and 20 mM for these assays and DCA alone had an IC50 of about 30 and 23 mM for M130429 and M010817, respectively (Suppl. Figure S 6e). Moreover, the glycolysis inhibitors 3-Bromopyruvate (3-BP) and 2-Desoxy-D-glucose (2DG) did not sensitize the cells to neocuproine (Suppl. Figure S6 f). These data indicate that the dynamic regulation of the PPP protects NRAS-mutated melanoma cells from oxidative stress and that the activity of the MAPK pathway is essential for this regulation.

**Figure 4:**
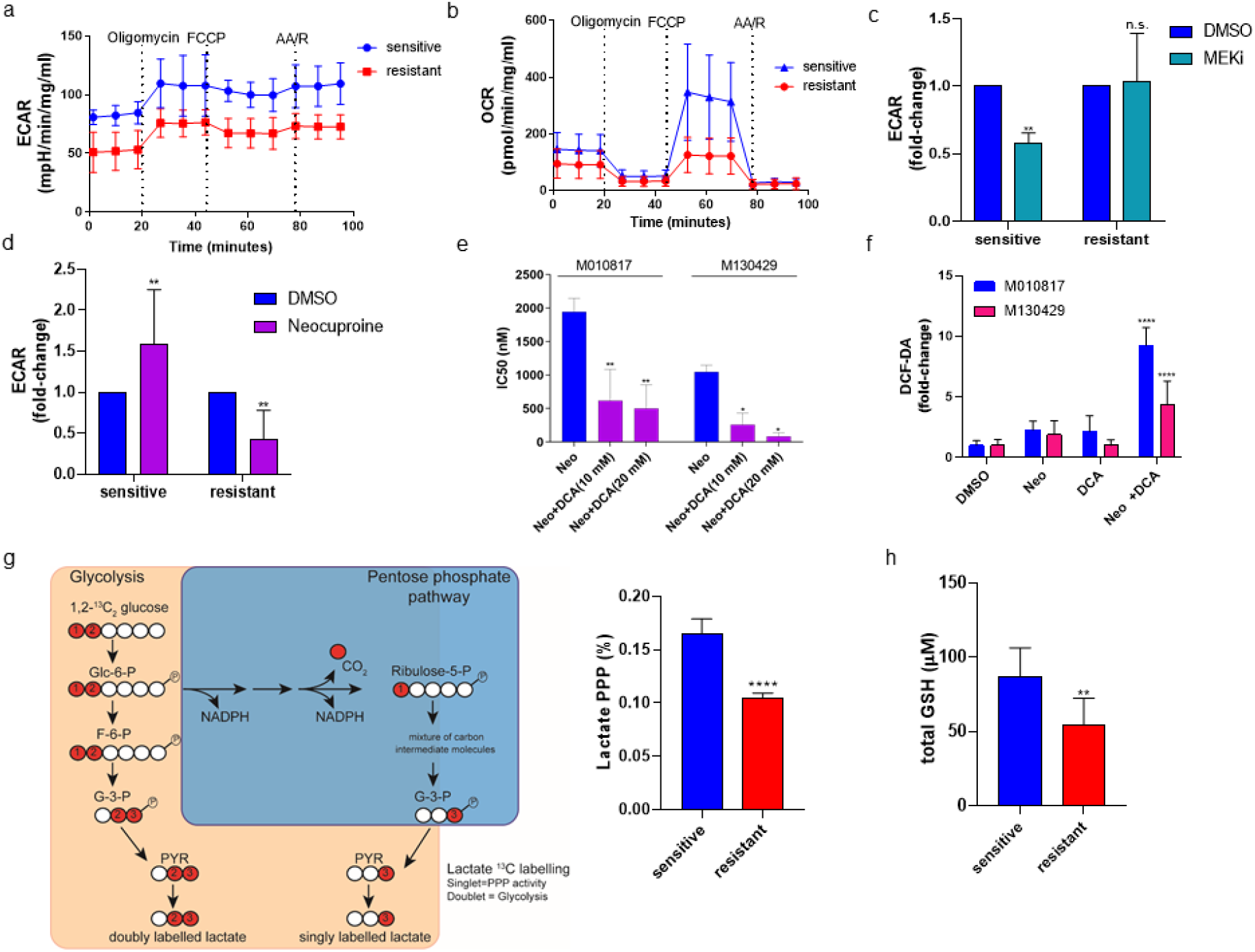
Low Glycolysis and PPP activity sensitizes resistant melanoma cells to ROS. We performed all following measurements with the cohort of two sensitive (M130429 and M010817) and four resistant (M130219, M130227, M121224, M160915) NRAS mutated melanoma cell lines (a) Extracellular acidification rate (ECAR) as a measurement of lactate production and (b) oxygen consumption rate (OCR) in living cells was measured by Agilent’s Seahorse technology. (c + d) Bar graphs represent the induction of ECAR (glycolytic reserve) in melanoma cells after treatment with MEKi (0.5 μM) or Neocuproine (1 μM) for 24 h (**p<0.01) (C) or (d) Neocuproine (**p<0.01), respectively. (e) IC50 values (growth inhibition) for Neocuproine of two MEKi sensitive melanoma cell lines alone or in the presence of DCA. (*p<0.05,**p<0.01, two-way ANOVA). (f) Two MEKi sensitive melanoma cell lines were treated with Neocuproine or the combination of Neocuproine and DCA. Induction of ROS was analyzed by FACS using DCF-DA (****p<0.0001, two-way ANOVA). (g) Graphical summary of how ^13^C-labled glucose is metabolized by glycolysis and PPP into differently labeled lactate. (h) MS analysis of the % of Lactate metabolized by the PPP in sensitive and resistant melanoma cell lines after the treatment with ^13^C-labled glucose (****p<0.0001, Student’s t-test). (h) Bar graph represents the total amount of Glutathione (GSH) in sensitive and resistant melanoma cells (**p<0.01, Student’s t-test).

ROS defense is achieved intracellularly through the GSH-NADPH axis. In the glycolysis pathway, NADPH has two different intracellular sources: PPP and TCA pathways through isocitrate dehydrogenase (IDH). In order to determine if the reduced PPP could increase basal ROS levels of resistant cells, we performed flux experiments with 13C-labled glucose to estimate the conversion of glucose molecules to lactate via glycolysis and PPP (Figure 4 g). Here, we found significantly reduced PPP activity in resistant melanoma cells, which could explain the lack of antioxidant defenses in resistant melanomas. Reduced GSH is considered one of the most important intracellular scavengers of ROS [52]. We also observed significantly reduced levels of total glutathione in our cohort of resistant melanoma cell lines (Figure 4 h).

### Neocuproine inhibits melanoma cell proliferation in patient-derived ex vivo tumors resistant to Binimetinib

Short-term cell culture systems are a powerful method for drug testing of primary tumor materials, especially because the cultures maintain some of their original stromal composition [53]. We collected fresh tumor material after surgery from three consenting melanoma patients with confirmed NRAS mutations and progressive tumors, following prior treatments (Suppl. Figure S8 a). Corresponding tumors were also kept in paraffin and stained for melanoma marker S100 and proliferation marker Ki67 to confirm viable melanoma tissue (Suppl. Figure S8 b). Fresh tumor material was sectioned using a vibratome in 400 μm slices and maintained in cell culture medium on cell culture inserts (Figure 5 a). They were treated with either MEKi (binimetinib), neocuproine, or a combination of both. *Ex vivo* slice cultures were incubated for an additional 72h under culture conditions, and immunohistochemistry was performed to evaluate tumor area and cell proliferation using S100 and Ki67 staining (Figure 5 b, Suppl. Figure S9). We found that two of the three patients were initially MEKi resistant, suggesting an intrinsic resistance mechanism; whereas, the third patient was partially but significantly sensitive to MEKi. In all cases, treatment with neocuproine significantly reduced the proliferation of tumor slice cultures, although no significant synergy was observed with the combination of MEKi. The lack of synergy between neocuproine and MEKi may be due to the lack of cell plasticity in this model and the short timing of 72h.

**Figure 5:**
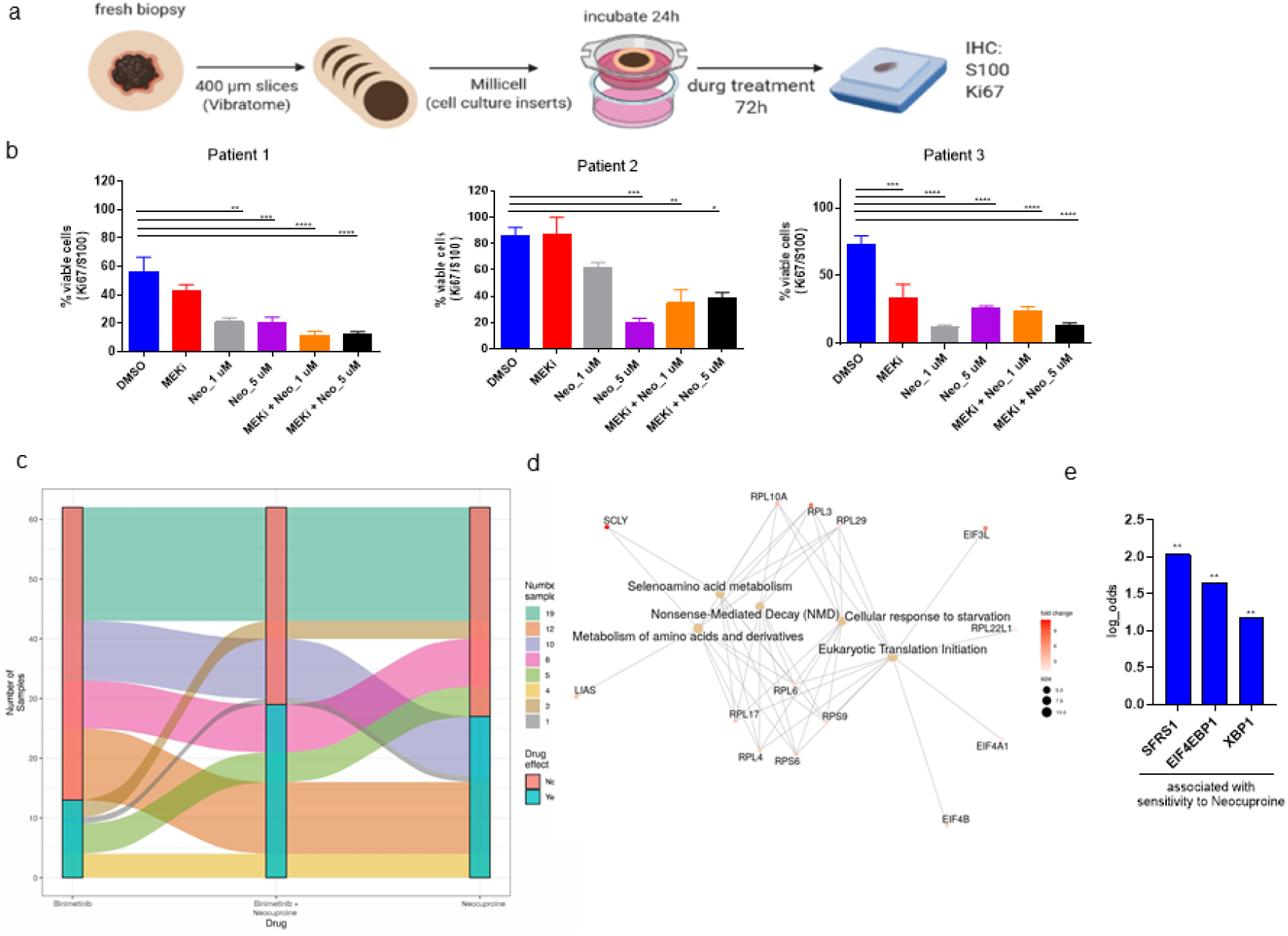
Neocuproine inhibits melanoma cell proliferation in patient-derived ex vivo orthologic slice-cultures. (a) Graphical summary showing the process of generating orthologic slice cultures (cartoon created using BioRender software). (b) Slice cultures were stained for S100 and Ki67 and the % of viable cells was calculated using QPath software (Suppl. Figure S9). The analysis of three melanoma patients is shown and % of viable cells is plotted as the mean of 5 individual measurement areas per tumor slice (**p<0.01, ***p<0.005, ****p<0.001, two-way ANOVA). (c) Alluvial-plot summarizing the responses to pharmacoscopy of 62 patient samples (TumorProfiler project). (d) Network analysis of mRNA biomarker (CCLE cell line screening) which correlated with sensitivity to Neocuproine (e) Biomarkers which corresponding with sensitivity to Neocuproine on the CCLE cell panel (FDR<0.01**).

Next, we validated neocuproine in a broader cohort of *ex vivo* clinical samples of the Tumor Profiler (TuPro) consortium. The TuPro study performed a multi-omics analysis in an approved clinical study (BASEC-2018-02050) in which ex vivo patient samples were assessed for their drug responsiveness employing different technologies beyond classical pathological and mutational analysis to guide treatment decisions [54]. Pharmacoscopy, a method to measure single-cell drug responses by immunofluorescence, automated microscopy, and image analysis, is part of the TuPro platform [55]. Neocuproine alone or in combination with binimetinib was assessed in tumor digests (Figure 5 c). A total of 62 melanomas were analyzed (Suppl. Table ST7). Of these, 49 (79%) were resistant to binimetinib. Most of the 49 tumor samples that were resistant to binimetinib were either sensitive to neocuproine monotherapy (10/49), combination with binimetinib (8/49), or both (12/49). Thus, the proportion of binimetinib failures that were sensitive to neocuproine monotherapy or in combination with binimetinib was significantly larger than those that were not (30/49 vs. 19/49, p = 0.023, 2-sample test for equality of proportions). Interestingly, 17 melanoma tumors were BRAF-mutated, highlighting the potential of neocuproine as a compound that can target resistant melanoma independent of their mutation status. In addition, part of the multi-level depiction of melanoma tumors was single-cell analysis using CyTOF technology on a panel of 53 pre-selected markers (Suppl. Figure S10 a). Using these data, we searched for proteins associated with the pharmacological response to neocuproine (Suppl. Figure S10 b). We found that a general downregulation of signaling pathways (e.g., MAPK or AKT) measured by their proteins and phosphorylated kinases was associated with sensitivity to neocuproine and binimetinib combination. To identify biomarkers distinguishing samples that responded to neocuproine and binimetinib combination, but not binimetinib alone, we identified the upregulation of CAV1, a protein that we recently reported to be associated with resistant and mesenchymal melanoma cells [24]. These data highlight that cells with low signaling activity but a mesenchymal phenotype can benefit from neocurpoine treatment alone or in combination with binimetinib. We validated these findings also on the original tumor material by immunohistochemistry (Suppl. Figure 10d). Clearly, the response to Binimetinib and neocuproine was associated with higher CAV1 staining but lower staining for pErk, pS6, or the melanocytic marker TYRP.

We recently demonstrated that phenotypic switching from melanocytic to mesenchymal melanoma sensitizes these cells to ferroptosis-inducing compounds [44]. In that study, cell lines with high mesenchymal scores were sensitive to ferroptosis-inducing compounds and dependent on GPX4. The significant relationship between melanoma cells with intrinsic high oxidative stress, mesenchymal phenotype, low cellular activity, and response to neocuproine prompted us to expand this observation to other cancers by taking advantage of the cell line screening platform at the Broad Institute (CCLE), which includes 486 cancer cell lines originating from different cancer entities [56]. Responses to neocuproine were observed in various cancer cell lines with no enrichment for a specific cancer type (Suppl. Figure S11 a-b, Suppl. Table ST8). This is not surprising given the heterogeneity of cancer cells. Therefore, we further analyzed the mRNA and protein biomarkers that were correlated with sensitivity to neocuproine (Figure 5 d-e). The mRNA network analysis resulted in pathway-enrichments in “Selenoamino-acid metabolism” or “cellular response to starvation” as well as “metabolism of amino acids and derivates” and “Eukaryotic translation initiation”. Selenoamino acid metabolism promotes oxidative stress by consuming NADPH and regulating the synthesis of selenoproteins such as TrxR1 and GPX4 [57, 58]. Cellular responses to starvation are associated with amino acid deprivation and reduced overall translation [58]. Moreover, we found that the positive expression of three protein biomarkers significantly correlated with sensitivity to neocuproine (SFRS1, EIF4EBP1 (non-phosphorylated), and XBP1), which are all involved in oxidative stress responses and translation depression [59–61].

### Combination therapy of neocuproine and binimetinib inhibits tumor growth and metastasis in vivo

To better understand the *in vivo* efficacy of neocuproine in combination with MEKi, we performed xenograft experiments using the NRAS-mutated and *in vitro* resistant melanoma cell line M160915. We found that MEKi and neocuproine were both effective in inhibiting M160915 tumor growth compared to tumor growth in vehicle-treated mice (neocuproine (p<0.005) and MEKi (p<0.0001) (Figure 6 a, Suppl. Figure S12 a). Nevertheless, the combination therapy showed significantly superior inhibitory effects compared to the single treatments (MEKi vs. Combi p<0.0003; neocuproine vs. combi p<0.0125), as well as an improved survival benefit (MEKi: p=0.037, hazard ratio (HR)=0.34; neocurpoine: p=0.31; HR=0.6; combined: p=0.001, HR=0.13).

**Figure 6:**
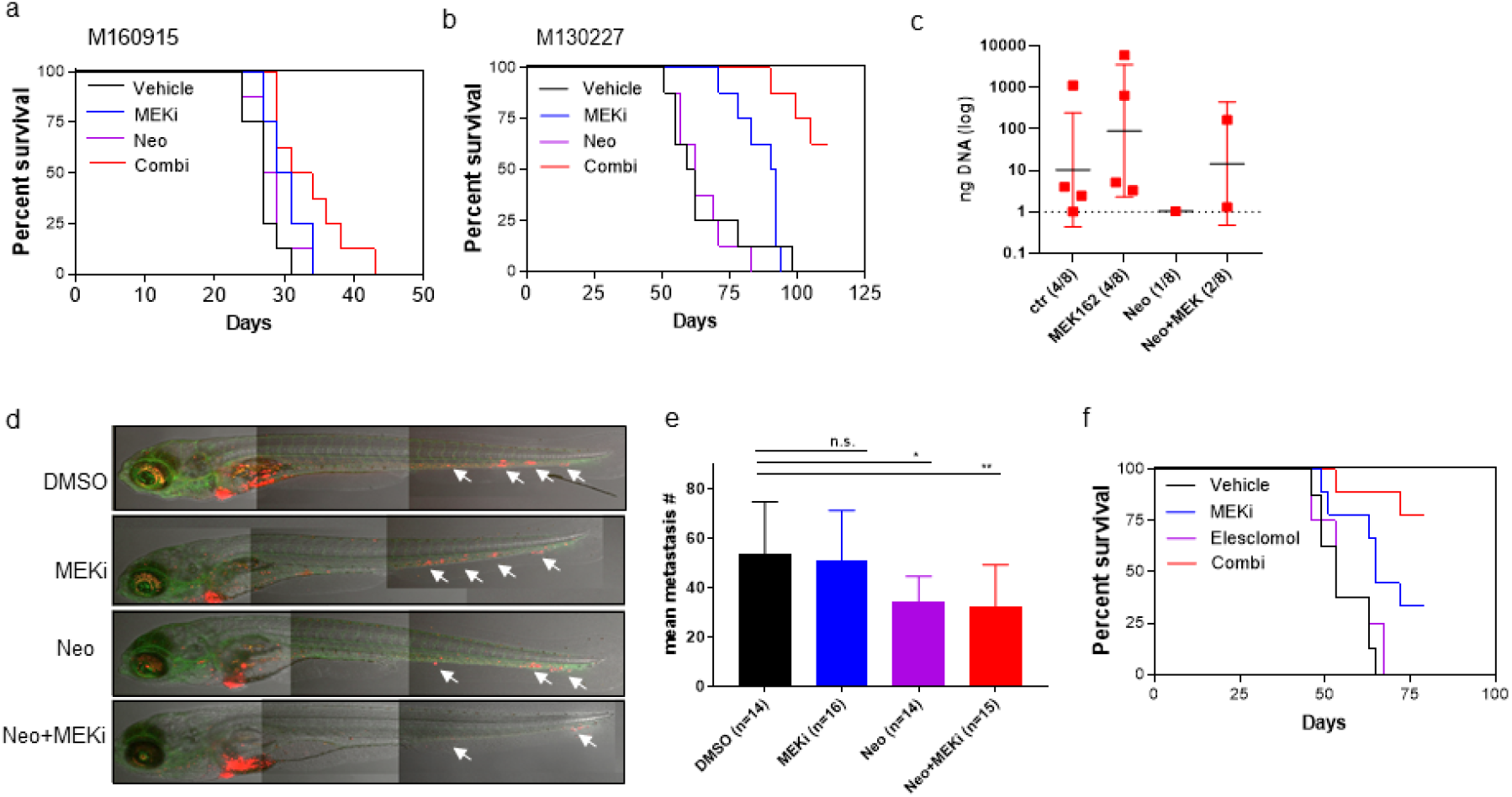
Combination therapy of Neocuproine and Binimetinib (MEK162) inhibits tumor growth and metastasis *in vivo*. (a) Kaplan-Meier-Survival curves of mice injected with M160915 melanoma cells and treated with MEKi (Binimetinib), Neocuproine (=Neo) or the combination (combi) (n=8) (b) Kaplan-Meier-Survival curves of mice injected with M130227 melanoma cells and treated with MEKi (Binimetinib), Neocuproine or the combination (combi) (n=8) (c) Quantification of the amount of human DNA in the draining lymph node using TaqMan PCR. Each dot represents the lymph node of one mouse with the amount of DNA (ng). (d) Images of Zebrafish larvae 4 days after the injection with red fluorescence labeled melanoma cells (M130227). Zebrafish are genetically engineered to express GFP in their entire vasculature under the control of the fli1 promoter. (e) Zebrafish have been treated with compounds for 72h before number of metastasis have been evaluated (*p<0.05, **p<0.01, two-way ANOVA). (f) Kaplan-Meier-Survival curves of mice injected with M130227 melanoma cells and treated with MEKi (Binimetinib), Elesclomol or the combination (combi) (n=8).

The major cause of death from melanoma is the spread of metastases through the blood, lymph, and visceral organs, which experience higher levels of oxidative stress than the established subcutaneous tumors in mice. Therefore, these cells are highly dependent on an intact antioxidant defense mechanism [62, 63]. The use of compounds that increase intracellular ROS, such as methotrexate, has been demonstrated to inhibit metastasis but not primary tumor growth. Here, we hypothesized that combining MEKi with neocuproine would have a significant impact on tumor growth and metastasis. We injected melanoma cells (M130227) that had metastasized to the draining lymph nodes. When tumors had grown palpable, mice were treated with vehicle alone, MEKi, neocuproine, or a combination of MEKi+ neocuproine (combi) (Suppl. Figure S12 b). We observed a statistically significant reduction in tumor volume when mice were treated with MEKi compared with the vehicle-treated group (p<0.0001) and the combination of MEKi and neocuproine compared with MEKi treatment alone (p<0.0001). No significant effect on tumor growth was observed in mice treated with neocuproine alone. Treatment with the combination of MEKi and neocuproine also significantly prolonged survival compared to vehicle treatment (MEKi: p=0.134, HR=0.45; neocuproine: p=0.27, HR=1.85; combi: p=0.001, HR=0.048) (Figure 6 b, Suppl. Figure S12 b) To examine metastasis formation, we removed the draining lymph nodes (DLN) and checked for the presence of human DNA using the Taqman PCR assay [46]. Four out of eight DLNs in the vehicle treatment and MEKi treated groups contained detectable human DNA; treatment with neocuproine or the MEKi-neocuproine combination resulted in DLNs without detectable human DNA (Figure 6 c). Although neocuproine did not inhibit tumor growth in the primary tumor, it did inhibit metastasis, suggesting that neocuproine targets invasive cell features that are sensitive to ROS induction. To investigate the specific effect of neocuproine on the inhibition of cancer cell dissemination, we used a zebrafish in vivo model. In this study, we injected fluorescently labeled M130227 cells into the yolk of 3-day-old zebrafish larvae following treatment with neocuproine, MEKi, or their combination (Figure 6 d-e). We performed whole-body imaging of zebrafish larvae and quantified the disseminated tumor cells. Melanoma cells in zebrafish remained resistant to MEK inhibitor treatment after addition of MEKi. Administration of neocuproine significantly reduced the number of disseminated tumor cells on its own, while MEKi did not, which is in line with our observations in mouse DLN. We observed no synergistic effects when administering a combination of neocuproine and MEKi, suggesting that the duration of the experiment (5 days) may be too short to induce phenotypic switching and heterogeneity, which would be essential for the benefit of the combination therapy.

Intra-lesion heterogeneity is a hallmark of melanoma and represents a major obstacle for the treatment with targeted therapies. Homogenous melanoma cell cultures can give rise to heterogeneous lesions that reflect the human situation; therefore, combination therapies are required to target all cancer cell phenotypes important for disease expansion [40, 64]. Therefore, we analyzed tumors from the M130227-mouse experiment to determine changes in intra-tumor heterogeneity using single-cell RNA sequencing (scRNA-seq). We found that the resistant cell line M130227 had a mesenchymal cell phenotype, which was also distinct from that of tumor cells. These cells did not express melanocytic markers, as revealed by scRNA sequencing, but rather mesenchymal marker genes, such as activin A (INHBA), N-cadherin (CDH2), or TGFβ (TGFBI) (Figure S13 a). This is in line with the differential expression of the bulk RNA-seq data shown in Figure 3 a. This cell line correlates with the invasive/mesenchymal phenotype. After injection, the homogeneous melanoma cells resulted in heterogeneous melanoma tumors, with cells expressing either melanocytic or mesenchymal markers (Suppl. Figure S12 a and (b)). ScRNA sequencing analysis of fractions of melanocytic or mesenchymal markers among the treatment groups also showed that proliferating, vehicle-treated tumors had high melanocytic melanoma cell content. MEKi treatment significantly increased the proportion of melanocytic melanoma cells, suggesting a transient increase in melanocytic cells as a mechanism of resistance to MEK inhibition. Only the combination treatment of MEKi with neocuproine dramatically reduced the amount of proliferative melanocytic melanoma, suggesting that cells escaping the melanocytic state (from neocuproine treatment) regained sensitivity to MEK inhibition (Suppl. Figure S12 b). Marker heterogeneity was also confirmed on xenograft tumors by immunohistochemistry using cell phenotype markers MelanA and INHBA (Suppl Figure S12 d).

Our *in vivo* data suggest that targeting only one cellular phenotype in heterogeneous melanoma may not effectively inhibit tumor growth. Moreover, ROS inducers might have great clinical potential but are not suitable for use as single agents, especially in melanoma, in which combination with targeted therapy is important. In 2013, a clinical trial comparing paclitaxel alone or in combination with the ionophoric copper-chelator and ROS inducer elesclomol highlighted that elesclomol did not improve progression-free survival (PFS) [65]. Given our results with neocuproine, we treated melanoma tumors with either MEKi, elesclomol, or their combination and observed a statistically significant survival benefit with the combination therapy of elesclomol and binimetinib (p<0.001, HR 0.052) compared to binimetinib (p=0.024, HR=0.26) and elesclomol (p=0.531; HR=0.725) (Figure 6 f, Suppl. Fig. S12c). In line with the ionophoric copper-chelator neocuproine, depletion of Cu ions from the cell culture medium also prevented growth inhibition by elesclomol in melanoma cell cultures (Supp. Figure S12e). Disulfiram (DSF), a clinically approved drug for treating alcoholism by irreversible inhibition of the acetaldehyde dehydrogenase enzyme, has also been shown to have anticancer activities [66]. Its mechanism of action is believed to involve the conversion of DSF to diethyl dithiocarbamate (DDC). DDC is another ionophoric copper-chelator whose cytotoxic activity is dependent on Cu ions (Suppl. Figure 12f). Xenograft animals were treated with DSF, binimetinib, or a combination of both drugs. We found that binimetinib and disulfiram significantly inhibited tumor growth (p<0.0001), but the combination of both compounds was more effective than DSF treatment alone (DSF vs. Bini+DSF (p<0.0018); Bini vs. Bini+DSF (p<0.18). Unfortunately, the combination of DSF and binimetinib caused severe toxicity in mice (mainly intestinal inflammation); therefore, the study was limited in time. Although ionophoric copper-chelators share a common mode of action, they appear to have different safety profiles *in vivo*. Our studies suggest that the ionophoric copper-chelator neocuproine, together with binimetinib, could be a potent and safe treatment for MEK-resistant and heterogeneous melanomas.

## Discussion

Clinical management of melanomas with NRAS mutations is challenging. Although oncogenic mutations are well described and immunotherapies are an option, targeted therapies against this tumor entity are lacking. Heterogeneity is a major obstacle in melanoma treatment; while MEK inhibitors target melanocytic melanoma cells with NRAS mutations, mesenchymal cells do not respond to MEKi, suggesting their independence of MAPK signaling [39]. In general, melanoma tumors consist of a complex network of heterogonous cancer and stromal cells, and metabolic programs need to be adapted to meet the demand of uncontrolled growth and disease progression in these complex environments. One of the best studied examples of this metabolic adaptation is the Warburg effect were cancer cells increase glucose use in order to achieve rapid production of ATP through the conversion of glucose to lactate via the anabolic pentose phosphate pathway (PPP). The PPP is also essential for DNA/RNA synthesis as well as for NADPH recycling, which protects dividing cells from oxidative stress. This metabolic switch is supported by oncogenic mutations which accelerate the transition, but are also energy consuming. In order to escape the metabolic addiction to oncogenes (e.g. MAPK mutations) and withstand targeted therapies, cells can adapt their metabolism to other pathways. Others and we have described for BRAF mutated melanomas that resistance to MAPK inhibitors drive metabolic reprogramming and activation of OXPHOS or fatty acid oxidation (FAO). In complex heterogonous tumors like melanoma this is shaped by the microenvironment or the metastatic niche were neighboring cells like adipocytes or fibroblasts facilitate metabolic adaptation [15, 67, 68].

In contrast, we show that NRAS-mutated melanoma, when becoming resistant to MEK inhibitors, become metabolically “*silent*”. A recent proteomics analysis of a large cohort of tumors with respect to their response to immunotherapies also revealed an association between BRAF mutations and elevated levels of mitochondrial OXPHOS proteins; whereas, in BRAF-WT mutated melanomas (including NRAS mutations), metabolic protein upregulation was not observed [69]. In BRAF-mutated melanoma, targeting these altered metabolic pathways is possible by targeting elevated OXPHOS or FAO [13, 15]. For NRAS-mutated melanoma, we did not find a superior activated metabolic pathway associated with MAPK resistance, but we found that in the metabolic “*silent*” cellular phenotype, elevated ROS levels become the Achilles’ heel of resistant cells.

It was shown for metastasizing cells to largely depend on NADPH-recycling pathways (e.g., the folate pathway). Given the strong transcriptional correlation between resistance and elevated ROS levels in NRAS-mutated melanoma, it is logical that resistant melanoma upregulates PHGDH, an enzyme that metabolizes serine in the folate pathway and links MEKi resistance to melanoma metastasis [70]. It has recently been shown that successfully metastasizing melanomas suffer from high ROS levels owing to their low PPP [71]. Melanoma cell metabolic heterogeneity can give rise to efficient metastasis of cancer cells and the loss of the lactate transporter MCT1 plays a key role in jeopardizing the PPP pathway, resulting in high basal ROS levels. This is in line with the correlation between mesenchymal phenotype melanoma, low PPP, and high basal ROS levels described here.

It was recently reported that elesclomol, which is another ionophoric copper chelator, induces a special death program called “cuproptosis”, which is defined by caspase-independent apoptosis and sensitivity through targeting upregulated mitochondrial respiration rate. Importantly, and contrary to our results with neocuproine, this ROS inducing agent was not rescued by the NAC antioxidant. Unlike with cuproptosis, we have shown here that in NRAS-mutated melanoma cells, the mode of death with neocuproine is caspase-dependent and specifically targets cells with low mitochondrial respiration. Moreover, the effect is rescued by NAC, which suggests that neocuproine works through non-cuproptosis based mechanisms. Heterogeneity in melanoma and resistance to targeted therapy has been shown to be a dynamic process, which involves several transient phenotypes *in vivo* [42]. Xenograft tumors derived from a homogenous NRAS-mutated and MEKi -resistant melanoma culture led to heterogeneous tumors, as shown by scRNA-seq, where phenotypic switching allowed therapeutic escape. The strategy of cancer cells to survive as treatment-persistent residual tumor cells between the phase of partial response and relapse has recently been described, where tumor cells use an embryonic diapause-like adaption [72].

This “arrested development” results in a downregulation of metabolism (e.g., mitochondrial activity), protein synthesis, and proliferation, but on the other hand the authors observed upregulation of processes related to ECM reorganization and cell adhesion, processes also observed by us in the invasive/mesenchymal melanoma-phenotype. It was further shown that cancer cells can enter the persistent-cell stage upon transcriptional quiescence without the acquisition of new mutations or rare pre-existing clones to drive entry into the diapause-like state. It appears that mesenchymal and resistant melanoma cells enter a diapause-like cell state to confer resistance. With the acquisition of a “*silent*” metabolism, which seems to be beneficial for persistence, cells cannot recycle important detoxifying compounds such as NADPH and GSH. These factors are used by enzymes of the PPP or folate pathway, which subsequently slows down and eventually disappears. Here, we can link this cell state of persistence of NRAS-mutated melanoma closely with the upregulation of cytoplasmic ROS and use the altered cell metabolism for novel therapeutic combination therapy. We speculate that these “silent” melanoma cells use vesicle trafficking to passively uptake carbon sources and building blocks from the microenvironment and the surrounding stroma. One advantage is also that by this process, cancer cells can adapt to a new microenvironment by driving their metabolic program depending on the availability of extracellular nutrition. We found that CAV-1, a protein associated with caveolae formation and a lipid chaperone, was a predictive biomarker to the response of neocuproine in association with low signaling activity (pErk and pS6) and melanocytic marker expression (TYRP). CAV-1 was shown to be a prognostic marker also in other cancers and is associated with extracellular vesicle trafficking, endo- and exocytosis of lipids [73].

Interestingly, we found that the “*silent*” cell state could be further used as a biomarker for predicting the response to neocuproine. During the screening of the CCLE cell panel we observed that three biomarkers (XBP1, SRFS1, and EIF4EBP1) were not only indicators of cellular stress but also known to actively downregulate cellular activity, keeping a “*silent*” cell state in the times of crisis (e.g. during drug treatments). EIF4EBP1 is a transcription factor that is activated in endoplasmic reticulum (ER) stress in the unfolded protein response (UPR) in an attempt to defend against high intracellular ROS levels. Oxidative stress drives the dephosphorylation of EIF4EBP1, thereby inhibiting the formation of the initiation complex and decreasing overall protein translation in the cell [60]. SFRS1 is an mRNA-splice factor and binding protein in which chaperones untranslate RNA into stress granules during cellular stress, inhibiting the translation of “housekeeping” genes. Accumulation of cytoplasmic SFRS1 increases the formation of stress granules, contributing to translational depression [61]. A similar result was observed in our TuPro analysis of *ex vivo* patient biopsies, where significant correlations with the response to neocuproine/binimetinib were only seen in tumors with low signaling pathway activity and upregulation of the mesenchymal cell state marker Caveolin-1. Altogether, cellular silencing, close to the concept of “arrested development” described in the above mentioned publications, leads to higher intracellular ROS which can be used to target these cell phenotypes, and not only in melanoma.

By combining small molecules targeting melanocytic and mesenchymal melanoma cells simultaneously, we showed significant impairment of tumor growth and metastasis. Moreover, metastasizing melanoma cells, which have been shown to produce high ROS levels [62], were targeted by neocuproine alone, thereby avoiding the invasion of melanoma cells into the lymph node. Our data suggest that the combination of MEKi with small molecules, which induces intracellular ROS, targets melanoma, especially with NRAS mutations, thus putting clinical trials with ROS inducers (e.g., SYMMETRY trial using elesclomol) into a new light [65]. We have shown *in vivo* that neither a MEK inhibitor nor a ROS inducer alone sufficiently impairs the combination of tumor growth and metastasis, suggesting clinical trials that combine both compounds. This novel approach would put not only MEK inhibitors back on stage for melanoma patients with NRAS mutations, but also other compounds that would synergize with ROS inducers to target silent metabolic states that sensitize these tumor cells to ROS.

## Supporting information

Suppl tabel T1-T7

## Acknowledgments

The authors wish to thank Dr. Christian Stirnimann of NEXUS Personalized Health Technologies, ETH Zurich, Switzerland, for assistance with the small-molecule library screening.

The authors also thank Dr. Diana Behrens and Dr. Christian Rupp, EPO Berlin, for their assistance and performance in the M130227-Xenograft in vivo study.

The authors wish to thank the Functional Genomics Center of Zurich for their RNA-seq and HPLC analysis services.

This work was supported by the Swiss National Science Foundation (310030_149946) and Austrian Research Promotion Agency (FFG) project 7940628 (Danio4Can).

We thank Federica Sella and Mirka Schmid for technical support.

