## Supplementary material for "ROS induction as a strategy to target persister cancer cells with low metabolic activity in NRAS mutated melanoma": Suppl figure S1-S13

### Suppl Figure S1-13

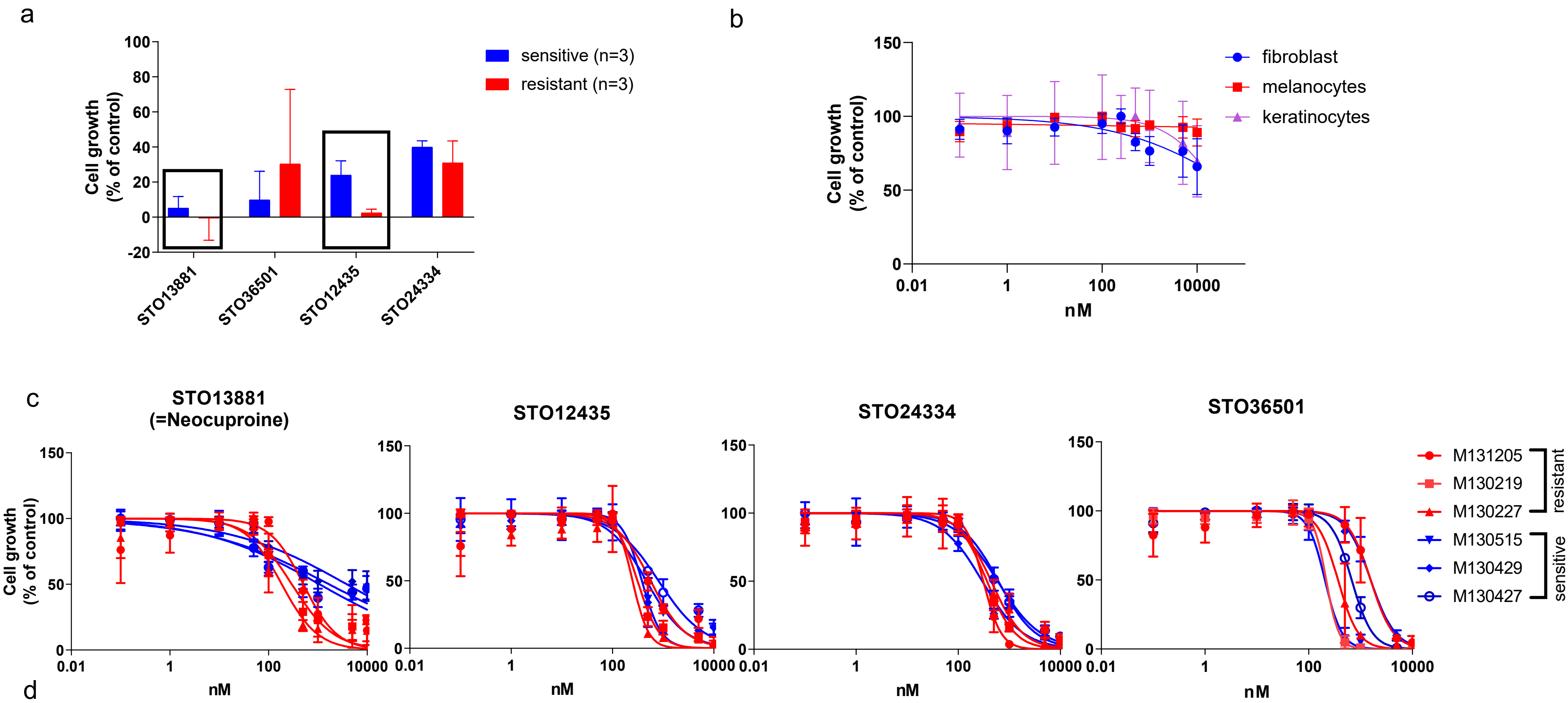


d

| | MEKi resistant | | | MEKi sensitive | | |
|---|---|---|---|---|---|---|
| | M131205 | M130219 | M130227 | M130515 | M130429 | M130427 |
| **STO13881** | | | | | | |
| IC50 | 511.8 | 297.3 | 185.9 | 1083 | 3877 | 1691 |
| 95% CI | 381 to 687.5 | 214.6 to 411.7 | 142.7 to 242.3 | 661.3 to 1775 | 2369 to 6348 | 848.4 to 3371 |
| **STO12435** | | | | | | |
| IC50 | 566.7 | 311.5 | 249.7 | 465.2 | 379.4 | 811.8 |
| 95% CI | 430.7 to 745.8 | 279.1 to 347.6 | 170.9 to 364.8 | 371.6 to 582.5 | 312.9 to 460.1 | 576.8 to 1143 |
| **STO24334** | | | | | | |
| IC50 | 310.3 | 365.7 | 442.3 | 612.2 | 296.9 | 592.2 |
| 95% CI | 245.4 to 392.3 | 305.4 to 437.8 | 381.5 to 512.9 | 560.4 to 668.8 | 256.6 to 343.4 | 466.9 to 751.3 |
| **STO36501** | | | | | | |
| IC50 | 1568 | 218.1 | 367.5 | 206 | 1569 | 676 |
| 95% CI | 1216 to 2022 | 184.3 to 258.1 | 381.5 to 512.9 | 182.3 to 232.8 | 1369 to 1799 | 619.0 to 739.5 |

Fig S1

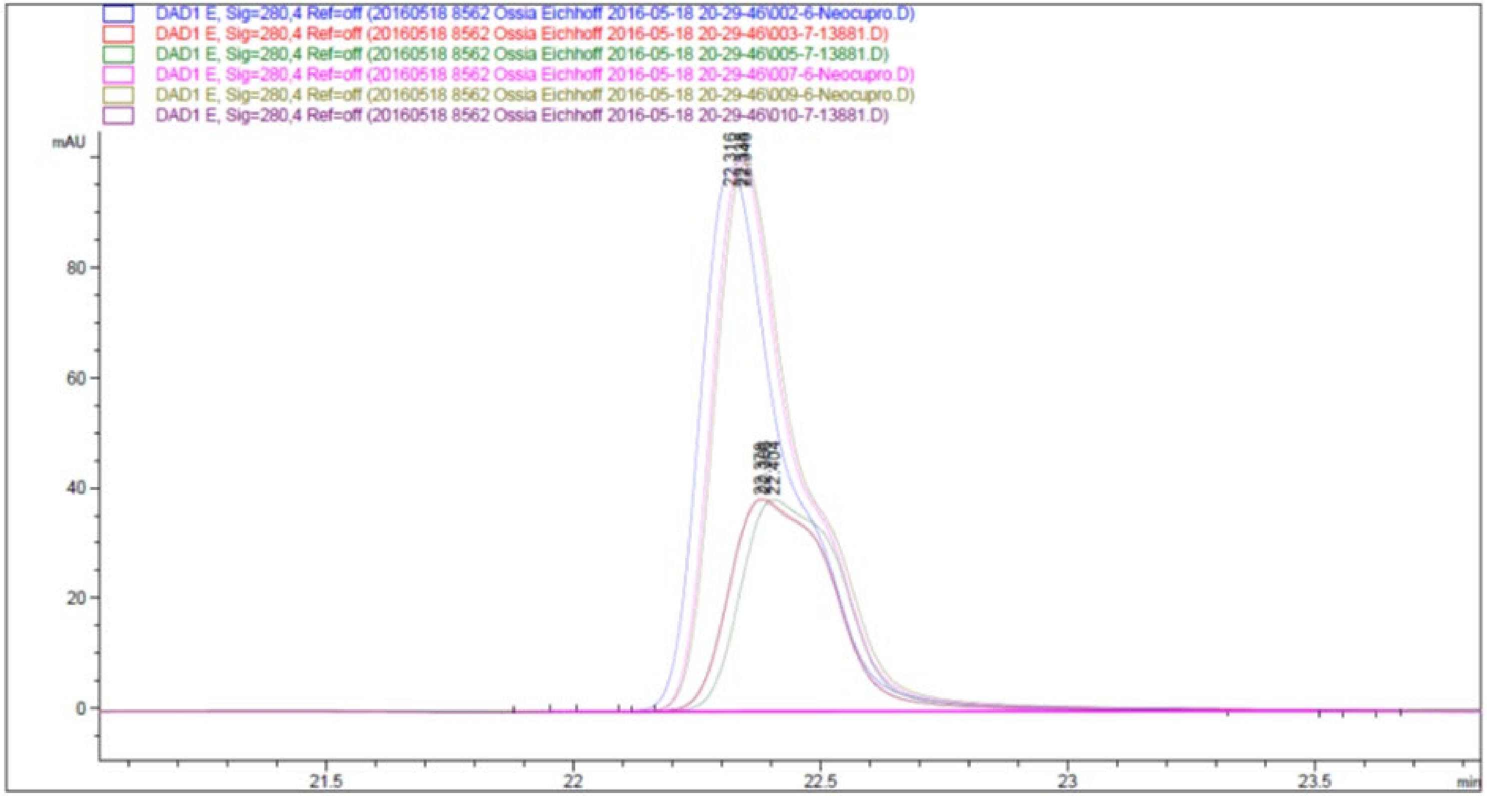
DAD1 E, Sig=280,4 Ref=off (20160518 8562 Ossia Eichhoff 2016-05-18 20-29-46\002-6-Neocupro.D)
DAD1 E, Sig=280,4 Ref=off (20160518 8562 Ossia Eichhoff 2016-05-18 20-29-46\003-7-13881.D)
DAD1 E, Sig=280,4 Ref=off (20160518 8562 Ossia Eichhoff 2016-05-18 20-29-46\005-7-13881.D)
DAD1 E, Sig=280,4 Ref=off (20160518 8562 Ossia Eichhoff 2016-05-18 20-29-46\007-6-Neocupro.D)
DAD1 E, Sig=280,4 Ref=off (20160518 8562 Ossia Eichhoff 2016-05-18 20-29-46\009-6-Neocupro.D)
DAD1 E, Sig=280,4 Ref=off (20160518 8562 Ossia Eichhoff 2016-05-18 20-29-46\010-7-13881.D)
mAU
22.316
80
60
40
20
0
21.5
22
22.5
23
23.5
min

Fig S2

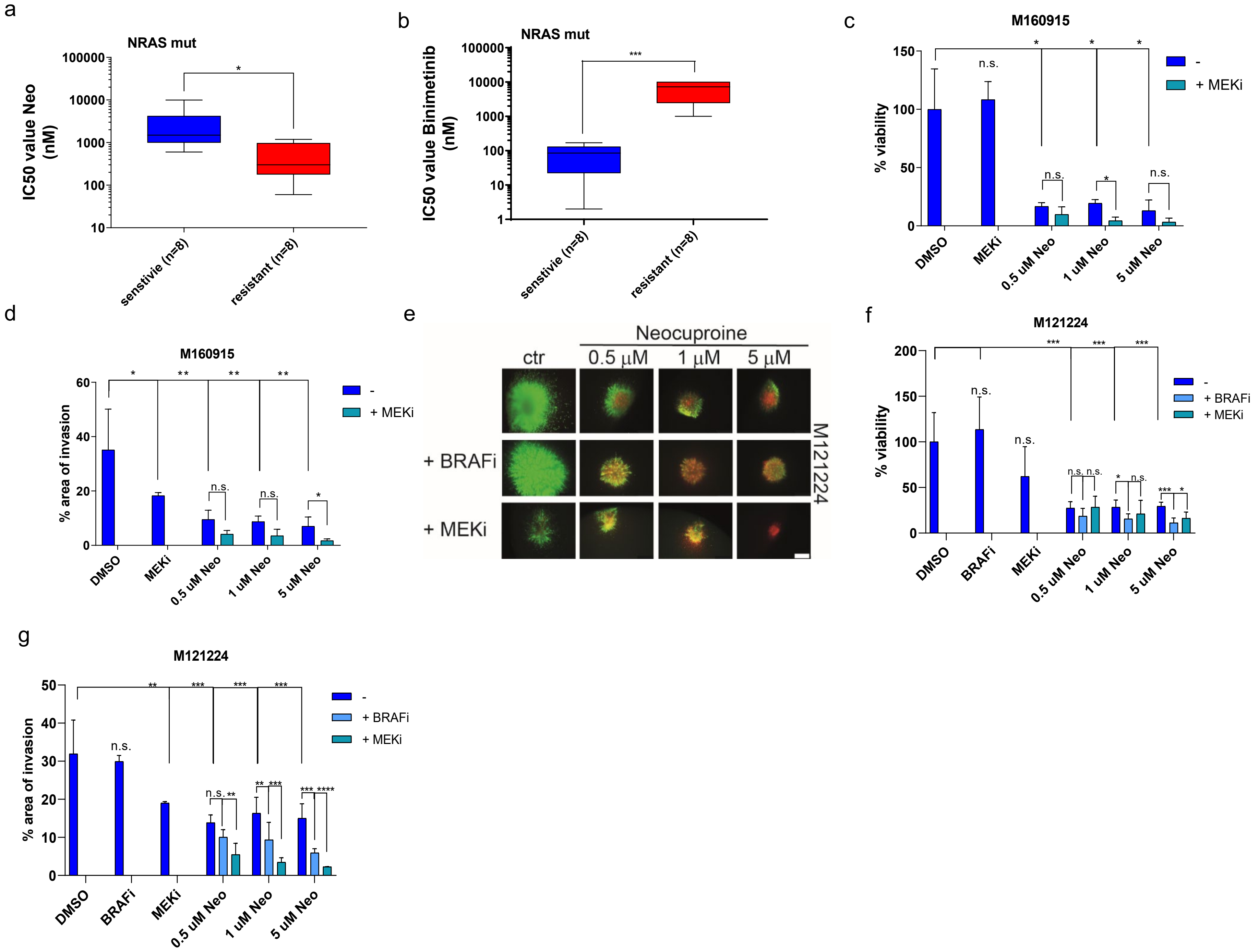
a
NRAS mut
IC50 value Neo (nM)
senstivie (n=8)
resistant (n=8)
b
NRAS mut
IC50 value Binimetinib (nM)
senstivie (n=8)
resistant (n=8)
c
M160915
% viability
- 
+ MEKi
DMSO
MEKi
0.5 uM Neo
1 uM Neo
5 uM Neo
d
M160915
% area of invasion
e
Neocuproine
ctr
0.5 µM
1 µM
5 µM
+ BRAFi
+ MEKi
M121224
f
M121224
% viability
+ BRAFi
BRAFi
g
M121224
% area of invasion

Fig S3

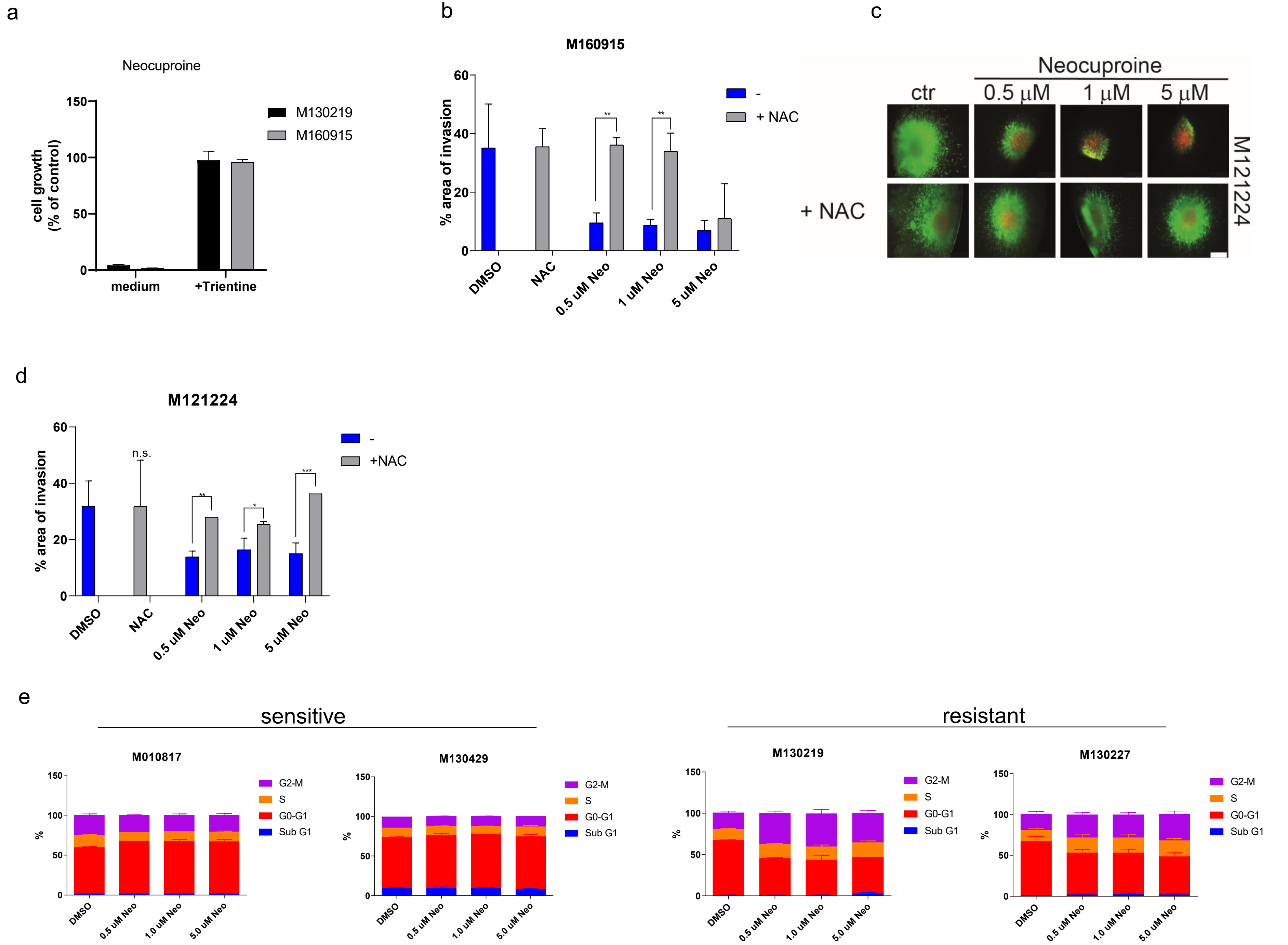
a
Neocuproine
cell growth (% of control)
150
100
50
0
M130219
M160915
medium
+Trientine
b
M160915
% area of invasion
60
40
20
0
-
+ NAC
**
**
DMSO
NAC
0.5 uM Neo
1 uM Neo
5 uM Neo
c
Neocuproine
ctr
0.5 μM
1 μM
5 μM
+ NAC
M121224
d
M121224
% area of invasion
60
40
20
0
n.s.
**
*
***
-
+NAC
DMSO
NAC
0.5 uM Neo
1 uM Neo
5 uM Neo
e
sensitive
resistant
M010817
M130429
M130219
M130227
%
150
100
50
0
G2-M
S
G0-G1
Sub G1
DMSO
0.5 uM Neo
1.0 uM Neo
5.0 uM Neo

Fig S4

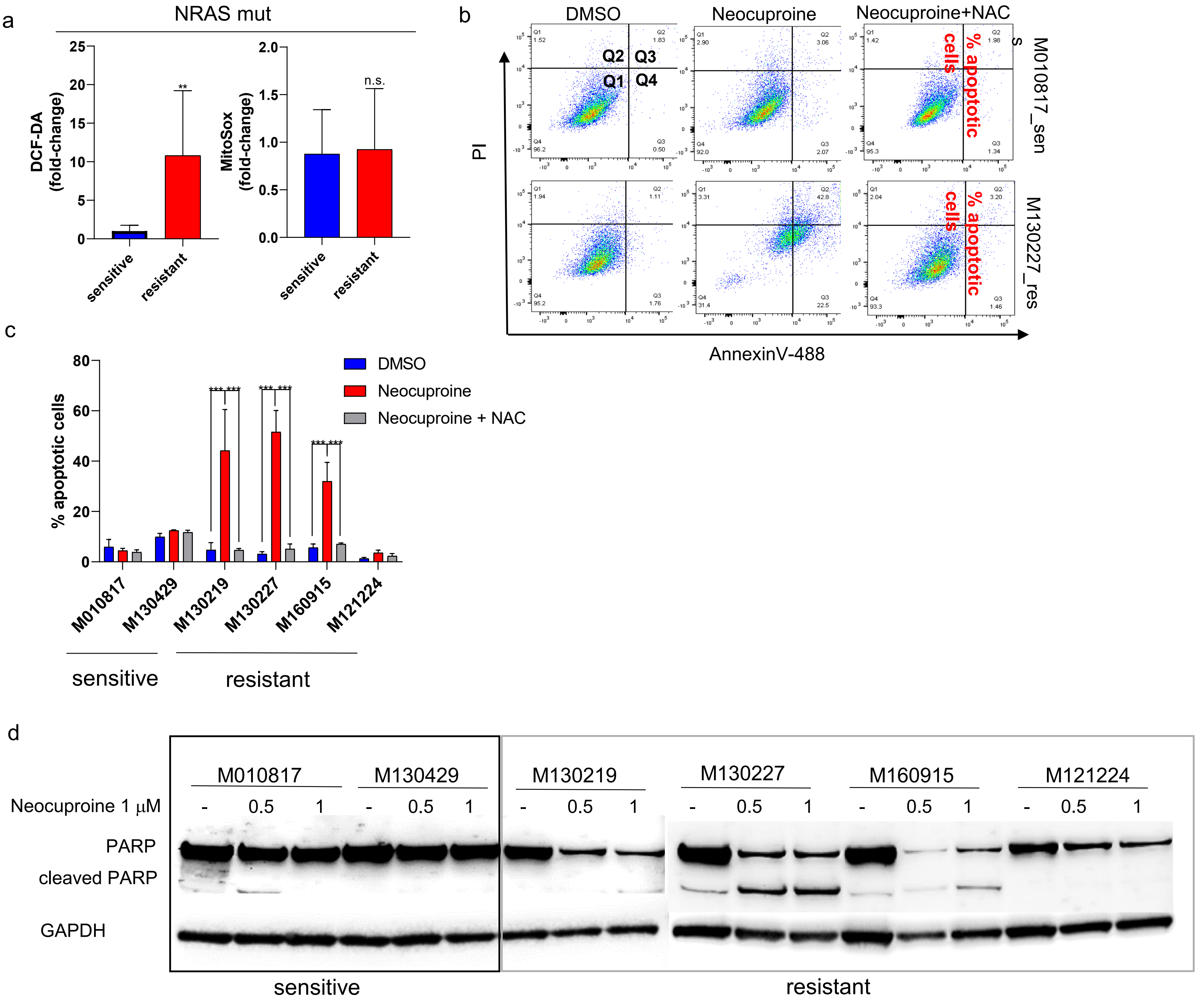
a
NRAS mut
DCF-DA (fold-change)
MitoSox (fold-change)
**
n.s.
sensitive
resistant
b
DMSO
Neocuproine
Neocuproine+NAC
Q1
Q2
Q3
Q4
% apoptotic cells
M010817_sen
M130227_res
PI
AnnexinV-488
c
% apoptotic cells
DMSO
Neocuproine
Neocuproine + NAC
M010817
M130429
M130219
M130227
M160915
M121224
sensitive
resistant
d
Neocuproine 1 μM
PARP
cleaved PARP
GAPDH
M010817
M130429
M130219
M130227
M160915
M121224
sensitive
resistant

Fig S5

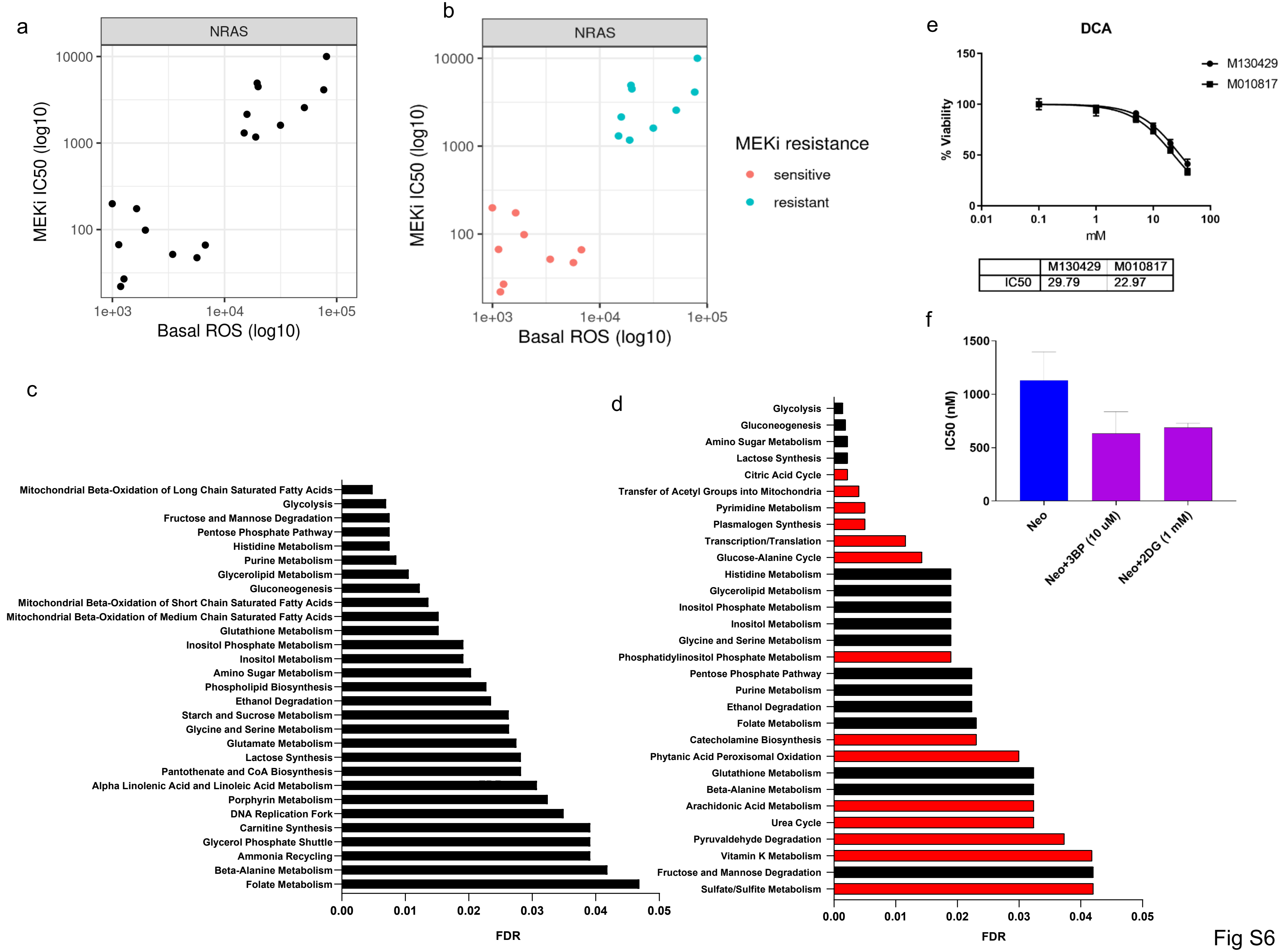
a
NRAS
MEKi IC50 (log10)
10000
1000
100
1e+03
1e+04
1e+05
Basal ROS (log10)
b
NRAS
MEKi IC50 (log10)
Basal ROS (log10)
MEKi resistance
sensitive
resistant
e
DCA
% Viability
150
100
50
0
0.01
0.1
1
10
100
mM
M130429
M010817
| M130429 | M010817
IC50 | 29.79 | 22.97
f
IC50 (nM)
1500
1000
500
0
Neo
Neo+3BP (10 uM)
Neo+2DG (1 mM)
c
Mitochondrial Beta-Oxidation of Long Chain Saturated Fatty Acids
Glycolysis
Fructose and Mannose Degradation
Pentose Phosphate Pathway
Histidine Metabolism
Purine Metabolism
Glycerolipid Metabolism
Gluconeogenesis
Mitochondrial Beta-Oxidation of Short Chain Saturated Fatty Acids
Mitochondrial Beta-Oxidation of Medium Chain Saturated Fatty Acids
Glutathione Metabolism
Inositol Phosphate Metabolism
Inositol Metabolism
Amino Sugar Metabolism
Phospholipid Biosynthesis
Ethanol Degradation
Starch and Sucrose Metabolism
Glycine and Serine Metabolism
Glutamate Metabolism
Lactose Synthesis
Pantothenate and CoA Biosynthesis
Alpha Linolenic Acid and Linoleic Acid Metabolism
Porphyrin Metabolism
DNA Replication Fork
Carnitine Synthesis
Glycerol Phosphate Shuttle
Ammonia Recycling
Beta-Alanine Metabolism
Folate Metabolism
0.00
0.01
0.02
0.03
0.04
0.05
FDR
d
Glycolysis
Gluconeogenesis
Amino Sugar Metabolism
Lactose Synthesis
Citric Acid Cycle
Transfer of Acetyl Groups into Mitochondria
Pyrimidine Metabolism
Plasmalogen Synthesis
Transcription/Translation
Glucose-Alanine Cycle
Histidine Metabolism
Glycerolipid Metabolism
Inositol Phosphate Metabolism
Inositol Metabolism
Glycine and Serine Metabolism
Phosphatidylinositol Phosphate Metabolism
Pentose Phosphate Pathway
Purine Metabolism
Ethanol Degradation
Folate Metabolism
Catecholamine Biosynthesis
Phytanic Acid Peroxisomal Oxidation
Glutathione Metabolism
Beta-Alanine Metabolism
Arachidonic Acid Metabolism
Urea Cycle
Pyruvaldehyde Degradation
Vitamin K Metabolism
Fructose and Mannose Degradation
Sulfate/Sulfite Metabolism
0.00
0.01
0.02
0.03
0.04
0.05
FDR

Fig S6

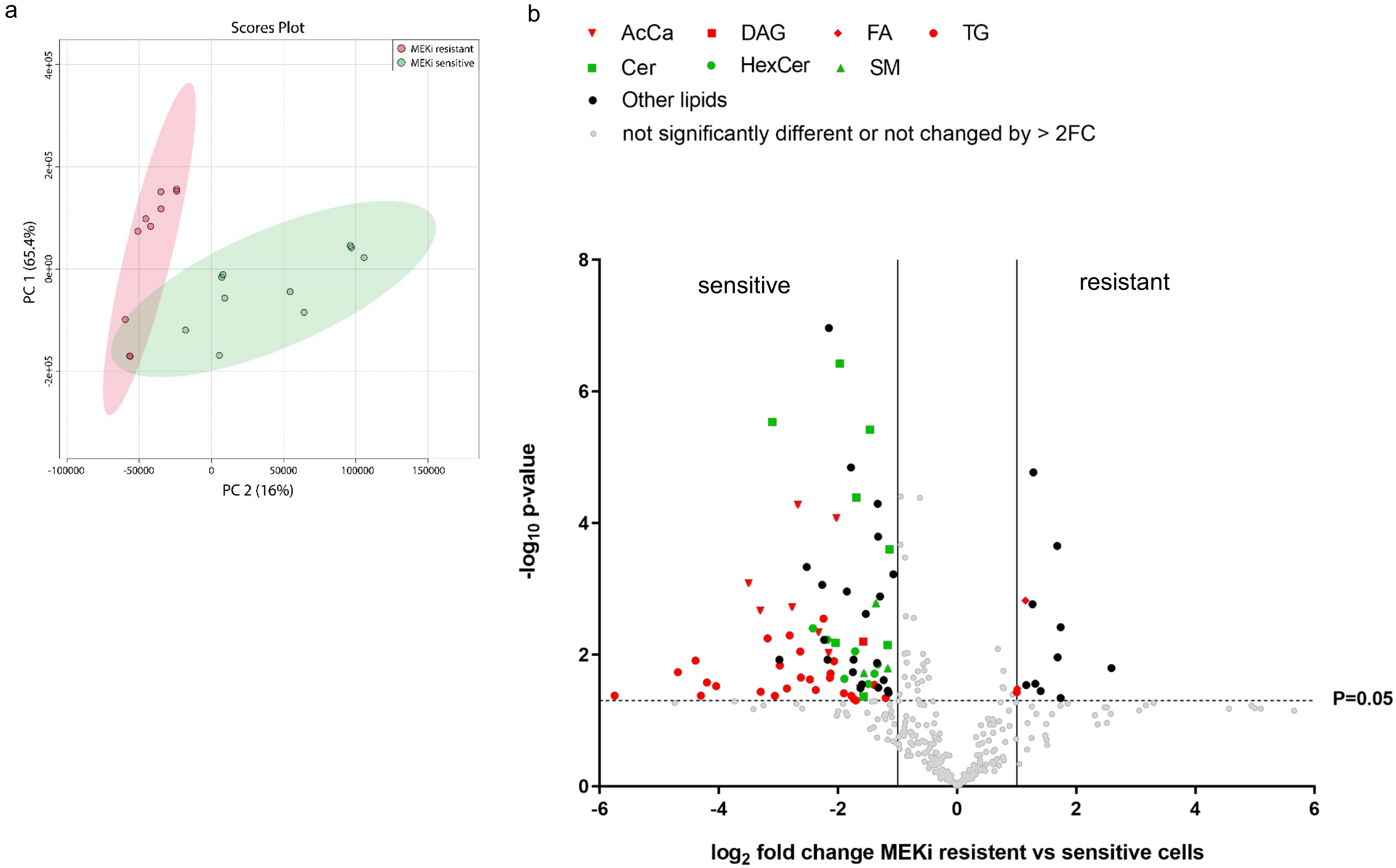
a
Scores Plot
MEKi resistant
MEKi sensitive
PC 1 (65.4%)
PC 2 (16%)
4e+05
2e+05
0e+00
-2e+05
-100000
-50000
0
50000
100000
150000
b
AcCa
DAG
FA
TG
Cer
HexCer
SM
Other lipids
not significantly different or not changed by > 2FC
sensitive
resistant
P=0.05
$-\log_{10}$ p-value
$\log_2$ fold change MEKi resistent vs sensitive cells
8
6
4
2
0
-6
-4
-2
0
2
4
6

Fig S7

a

| | description | confirmed mutation | treatment before surgery | ex vivo material |
|---|---|---|---|---|
| **Patient 1** | peritoneal metastasis | NRAS Q61R | ipilimumab (09/15-10/15),<br>Pembrolizumab (11/15-03/16): PD,<br>Dacarbazine (05/16-05/16): PD,<br>vindesin+cisplatin (07/16-10/16): PD,<br>taxol (02/17-07/17): mixed response | Aug 17 |
| **Patient 2** | small intestine metastasis | NRAS Q61K | Pembrolizumab+/-Epacadostat (03/16-03/17): PD;<br>Ipilimumab (04/17-07/17):PD | Okt 17 |
| **Patient 3** | inguinal lymph node metastasis | NRAS Q61K | ipilimumab+nivolumab (05/17-07/17):PD,<br>electrochemotherapy (05/17-07/17):PD | Okt 17 |

b

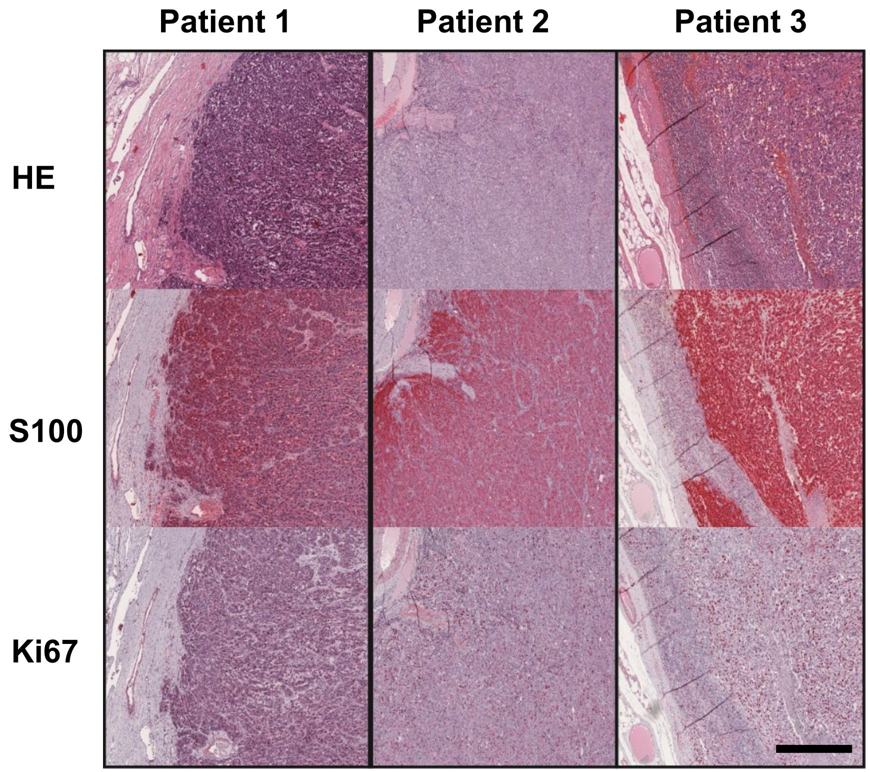


Fig SF8

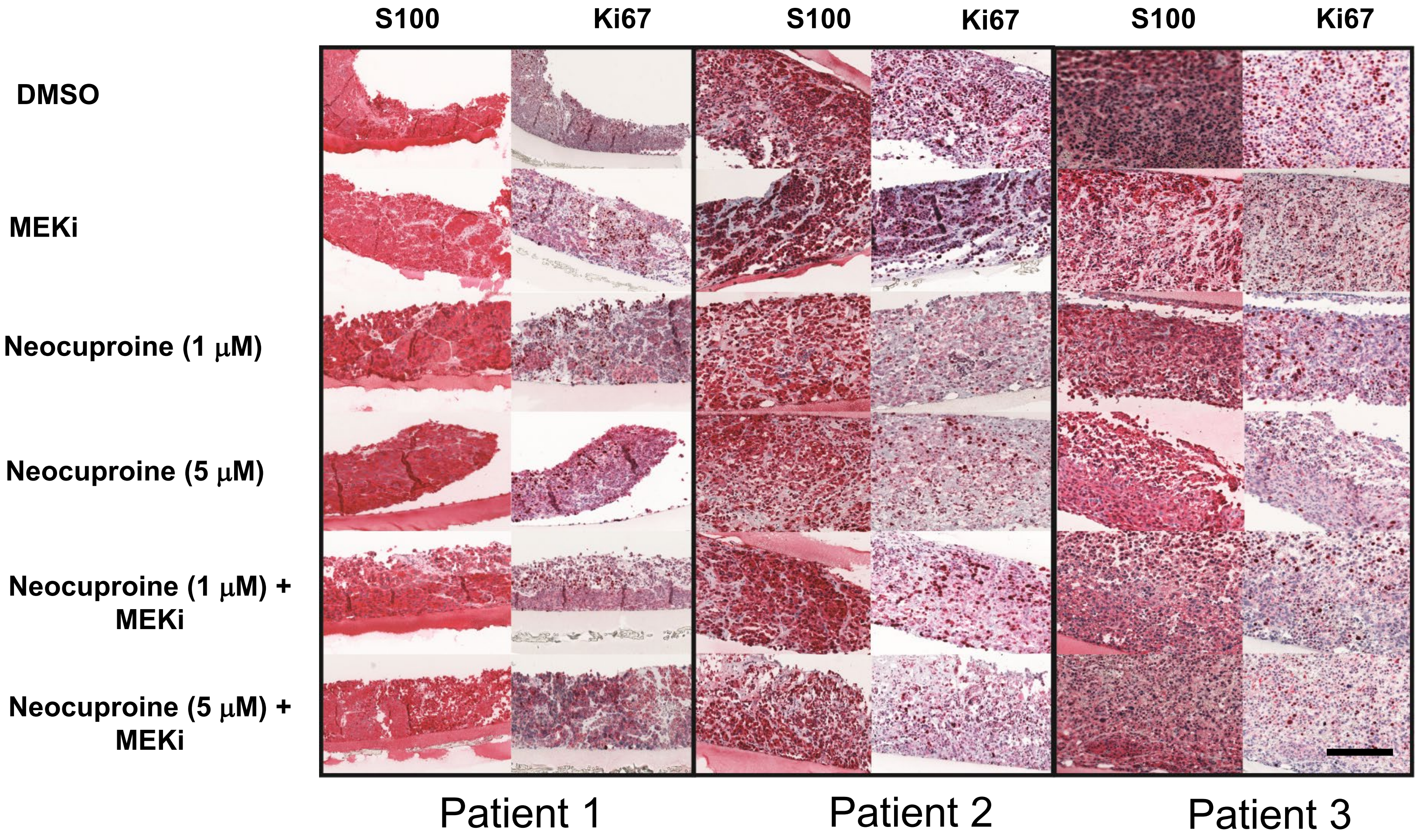
S100
Ki67
S100
Ki67
S100
Ki67
DMSO
MEKi
Neocuproine (1 μM)
Neocuproine (5 μM)
Neocuproine (1 μM) + MEKi
Neocuproine (5 μM) + MEKi
Patient 1
Patient 2
Patient 3

Fig S9

a

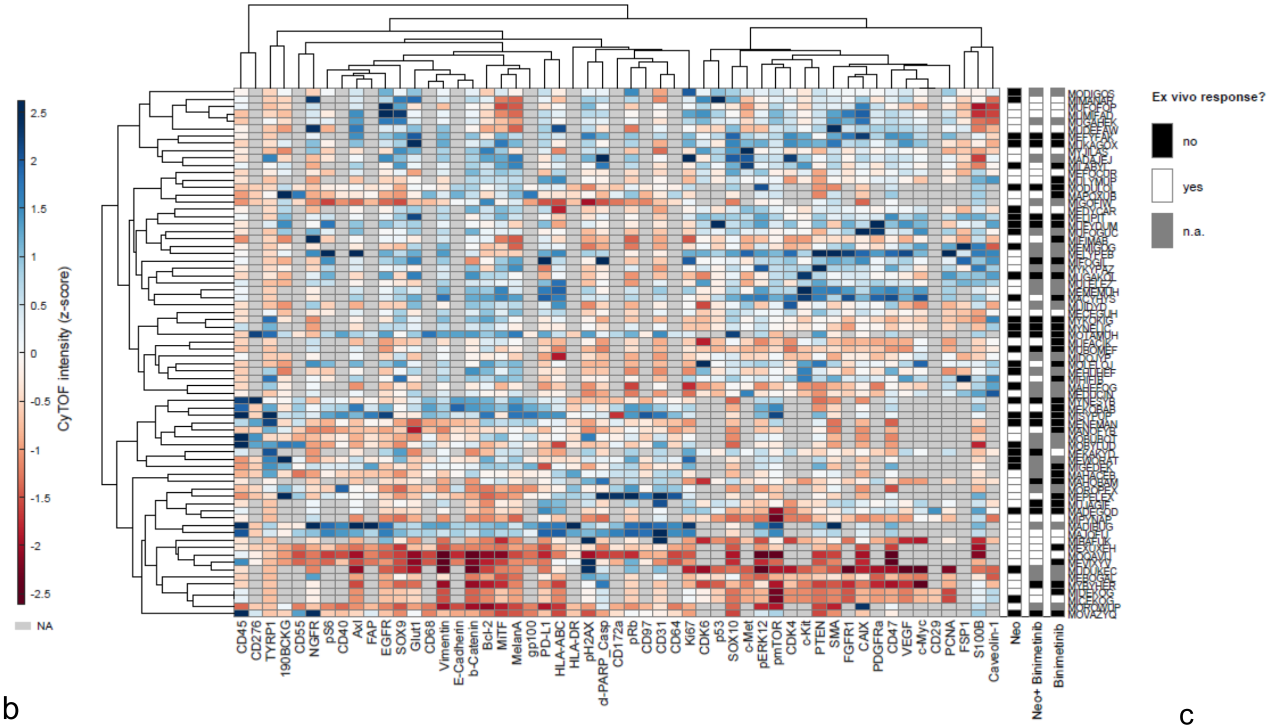
CyTOF intensity (z-score)
2.5
2
1.5
1
0.5
0
-0.5
-1
-1.5
-2
-2.5
NA
Ex vivo response?
no
yes
n.a.
CD45
CD276
TYRP1
190BCKG
CD55
NGFR
pS6
CD40
Axl
FAP
EGFR
SOX9
Glut1
CD68
Vimentin
E-Cadherin
b-Catenin
Bcl-2
MITF
MelanA
gp100
PD-L1
HLA-ABC
HLA-DR
pH2AX
d-PARP_Casp
CD172a
pRb
CD97
CD31
CD64
Ki67
CDK6
p53
SOX10
c-Met
pERK12
pmTOR
CDK4
c-Kit
PTEN
SMA
FGFR1
CAIX
PDGFRa
CD47
VEGF
c-Myc
CD29
PCNA
FSP1
S100B
Caveolin-1
Neo
Neo+ Binimetinib
Binimetinib

b

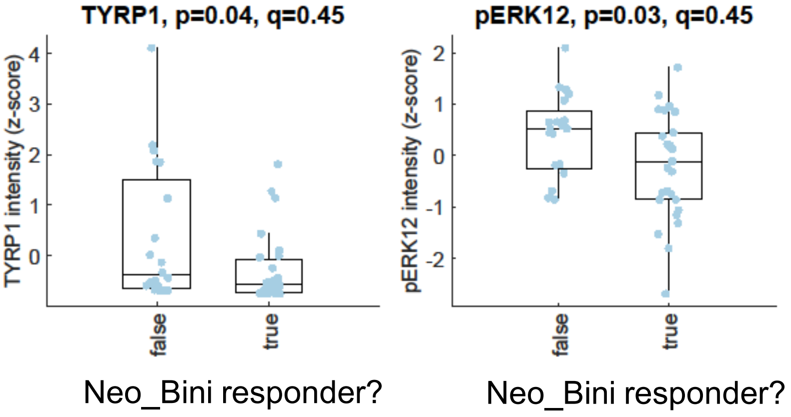
TYRP1, p=0.04, q=0.45
TYRP1 intensity (z-score)
false
true
Neo_Bini responder?
pERK12, p=0.03, q=0.45
pERK12 intensity (z-score)
false
true
Neo_Bini responder?
pS6, p=0.03, q=0.45
pS6 intensity (z-score)
false
true
Neo_Bini responder?

c

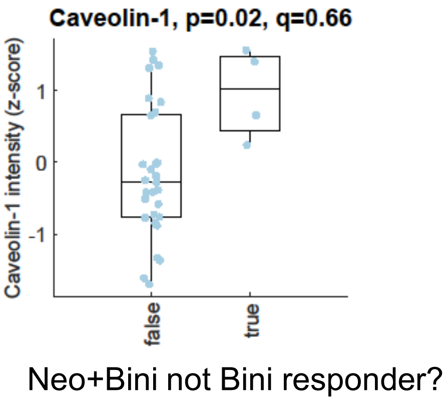
Caveolin-1, p=0.02, q=0.66
Caveolin-1 intensity (z-score)
false
true
Neo+Bini not Bini responder?

Fig S10

d

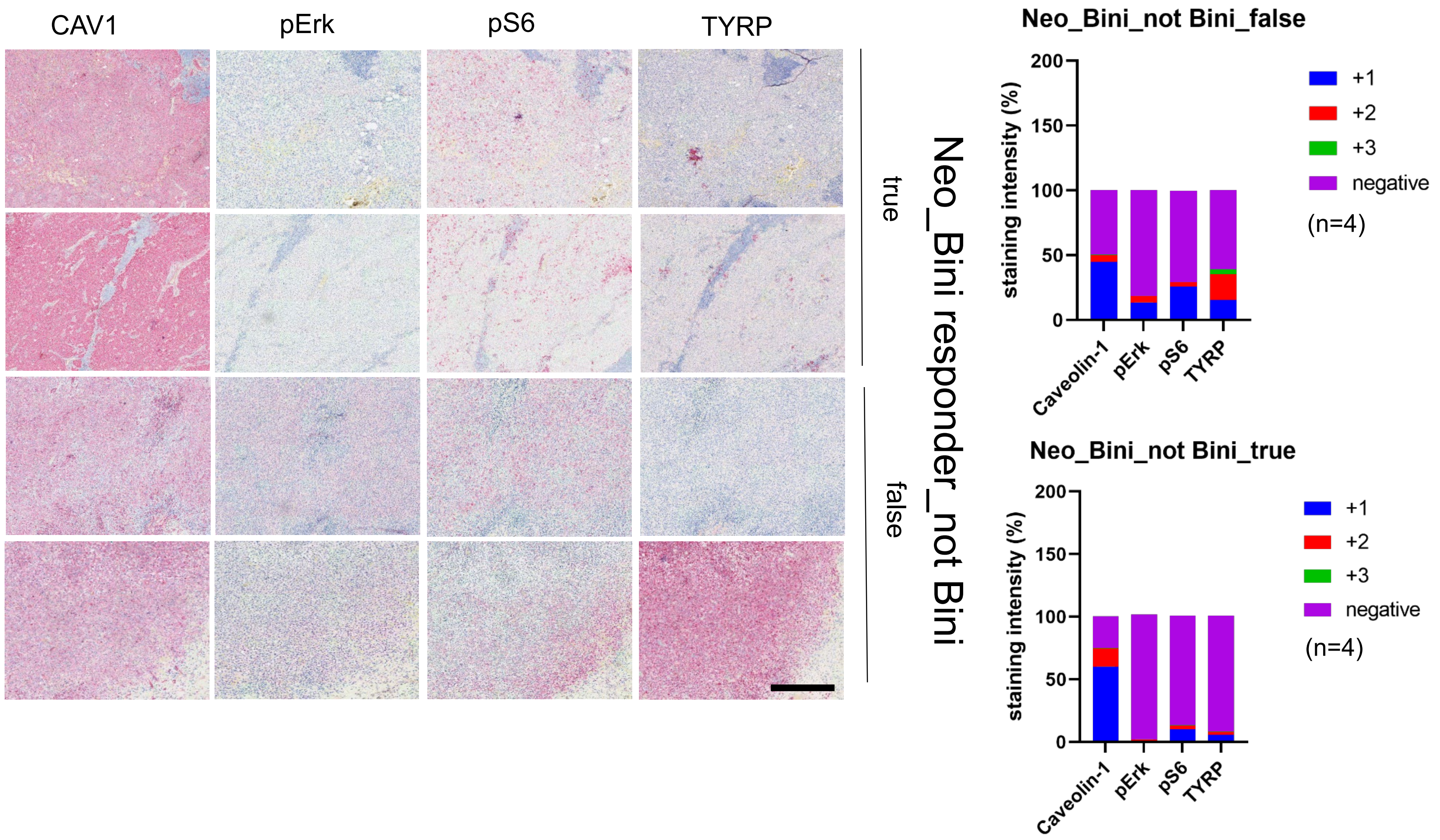
CAV1
pErk
pS6
TYRP
true
false
Neo_Bini responder_not Bini
Neo_Bini_not Bini_false
staining intensity (%)
200
150
100
50
0
Caveolin-1
pErk
pS6
TYRP
+1
+2
+3
negative
(n=4)
Neo_Bini_not Bini_true
staining intensity (%)
200
150
100
50
0
Caveolin-1
pErk
pS6
TYRP
+1
+2
+3
negative
(n=4)

Fig S10

a

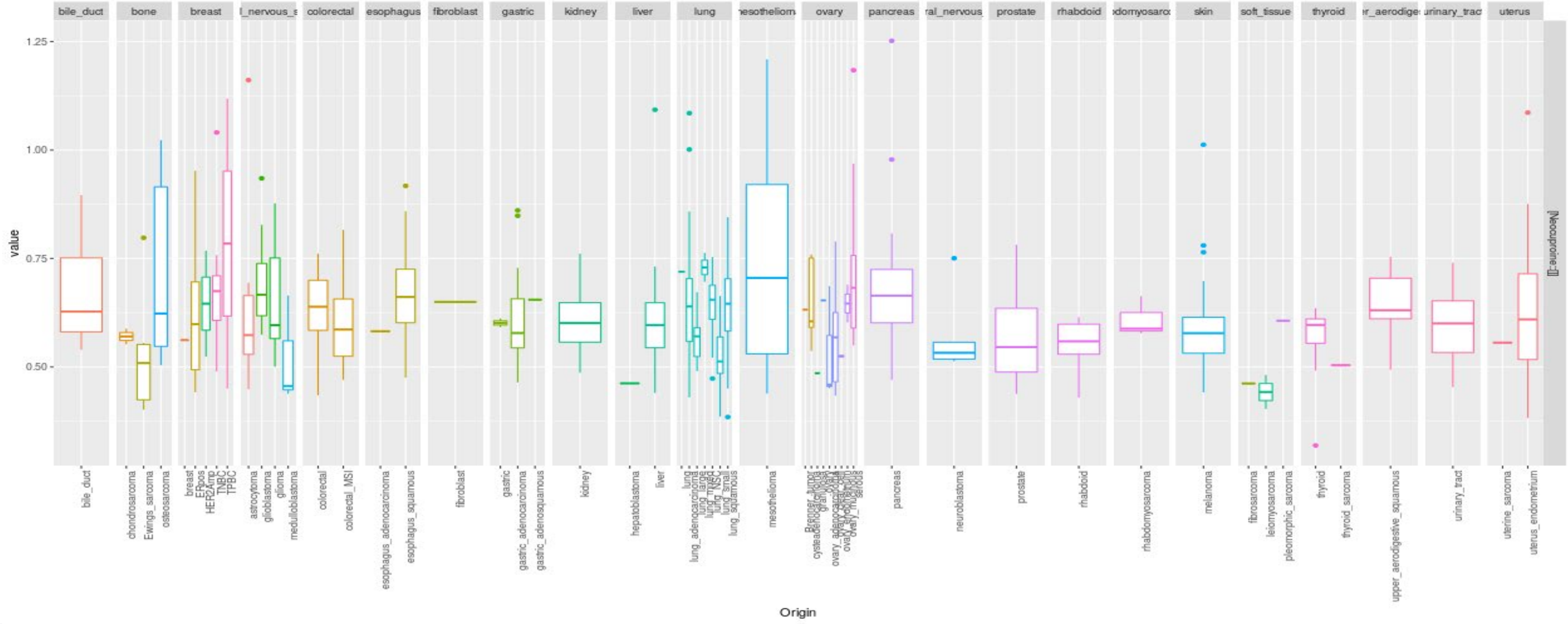

value
Origin
Neucuproine]


b

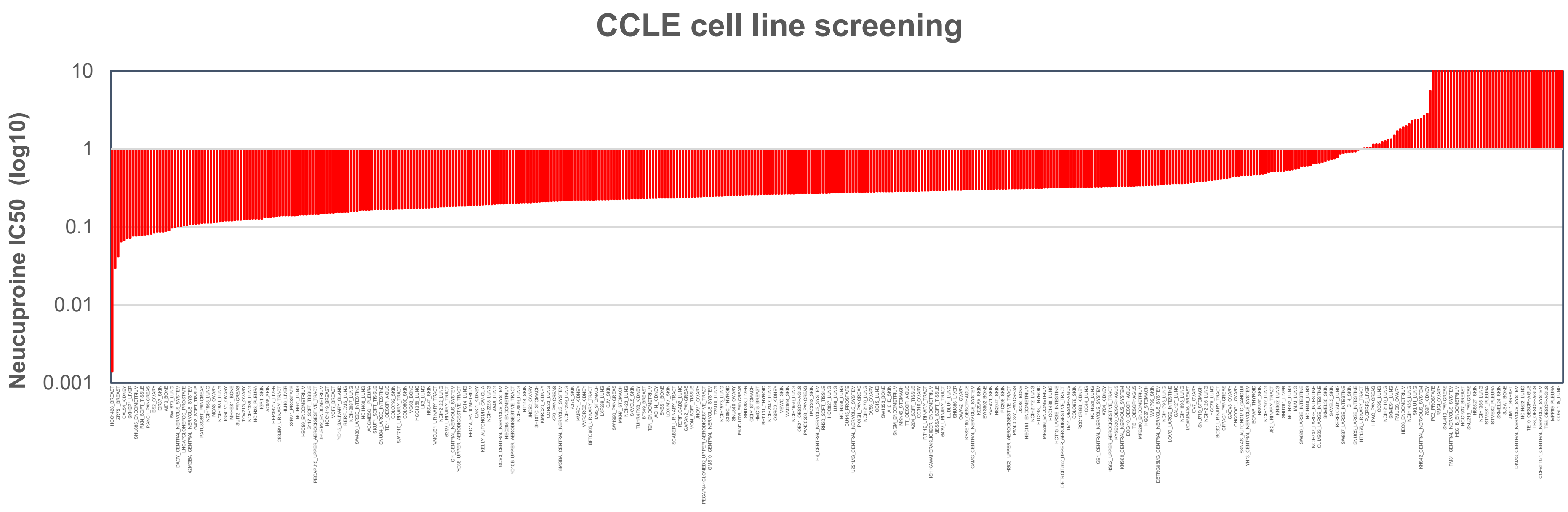

CCLE cell line screening
Neucuproine IC50 (log10)
10
1
0.1
0.01
0.001


Fig S11

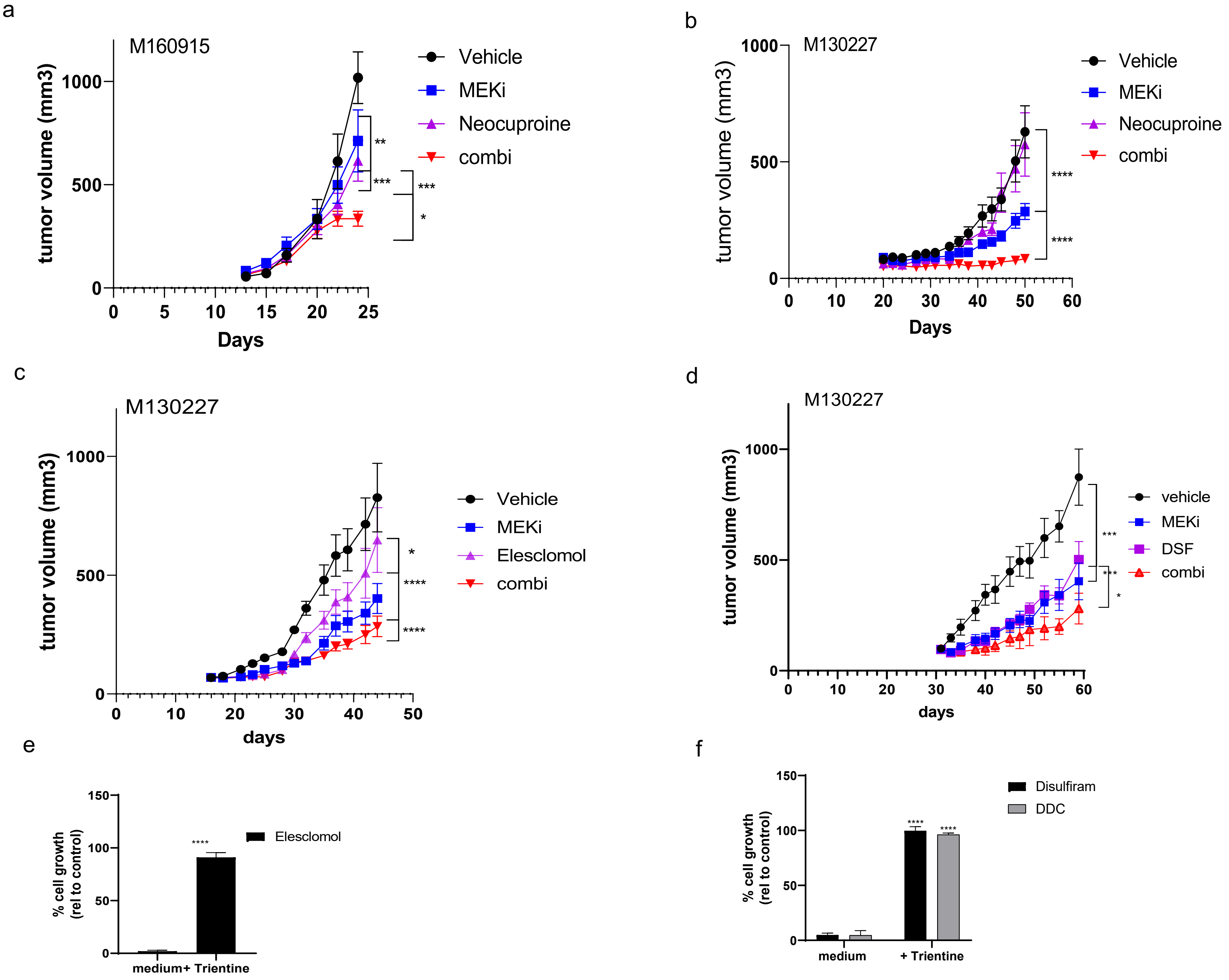
a
M160915
tumor volume (mm3)
Days
Vehicle
MEKi
Neocuproine
combi
b
M130227
tumor volume (mm3)
Days
Vehicle
MEKi
Neocuproine
combi
c
M130227
tumor volume (mm3)
days
Vehicle
MEKi
Elesclomol
combi
d
M130227
tumor volume (mm3)
days
vehicle
MEKi
DSF
combi
e
% cell growth (rel to control)
medium+ Trientine
Elesclomol
f
% cell growth (rel to control)
medium
+ Trientine
Disulfiram
DDC

Fig S12

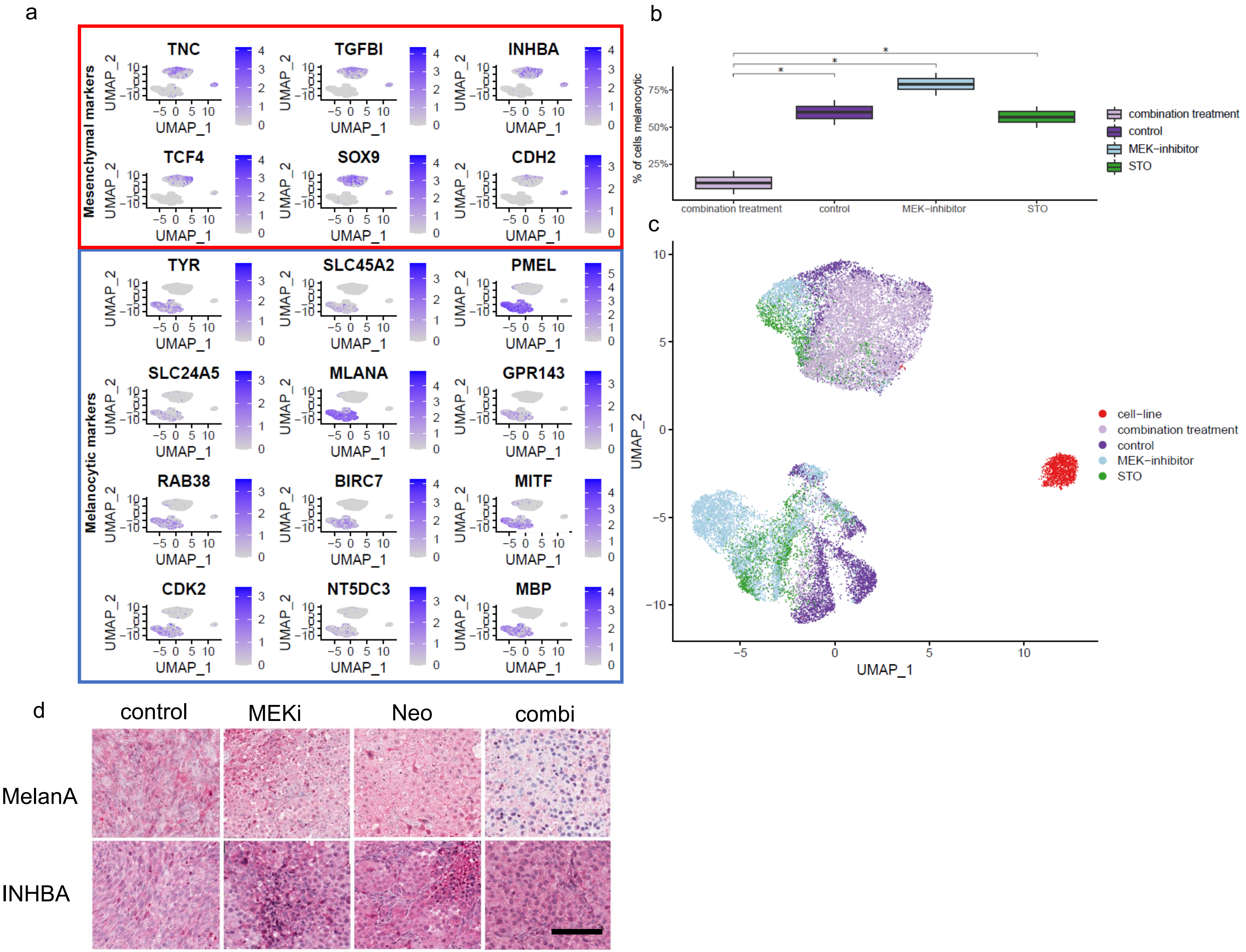
a
Mesenchymal markers
TNC
TGFBI
INHBA
TCF4
SOX9
CDH2
Melanocytic markers
TYR
SLC45A2
PMEL
SLC24A5
MLANA
GPR143
RAB38
BIRC7
MITF
CDK2
NT5DC3
MBP
UMAP_1
UMAP_2
b
% of cells melanocytic
combination treatment
control
MEK−inhibitor
STO
c
cell−line
combination treatment
control
MEK−inhibitor
STO
UMAP_1
UMAP_2
d
control
MEKi
Neo
combi
MelanA
INHBA

Fig S13
